# Comprehensive Functional Self-Antigen Screening to Assess Cross-Reactivity in a Promiscuous Engineered T-cell Receptor

**DOI:** 10.1101/2025.05.13.653646

**Authors:** Govinda Sharma, Fei Teng, James Round, Sophie A. Sneddon, Scott D. Brown, Sarania Sivasothy, Robert A. Holt

**Author notes:** Corresponding Author – Robert A. Holt, 7.114, 675 W 10^th^ Ave, Vancouver, BC, Canada V5Z 1L3.

## Abstract

T cell receptor therapeutics are an emerging modality of biologic and cell-based medicines with the unique ability to target intracellular antigens and finely discriminate between healthy and infected or mutated cells. An obstacle to the development of new T cell receptor therapeutics is the difficulty in engineering these proteins for enhanced therapeutic efficacy while avoiding introduction of unexpected off-target autoreactivity. In this study, we apply a functional high-throughput screening assay, Tope-seq, to detecting cross-reactive epitopes in libraries of >5 x 10^5^ unique peptide-coding sequences. We retrospectively analyze an affinity-enhanced engineered T cell receptor, which previously failed clinical trials due to severe off-target toxicity caused by epitope cross-reactivity, by comprehensive functional testing against all genome-coded self-antigens. Using the Tope-seq methodology, we were able to identify the epitope mediating off-target reactivity at a significance threshold of p < 0.01 in first-pass bulk screening. We also identified other potential cross-reactive epitopes of the engineered TCR of-interest, suggesting that the need for assessing promiscuity in TCR based therapeutics is larger than previously appreciated.

## Introduction

T cells are a critical piece of the vertebrate adaptive immune system responsible for removing pathogens and mutated cells from the body. Central to the function of T cells is the T cell receptor (TCR), a membrane-anchored immunoglobulin family member protein generated by the programmed somatic rearrangement of germline-encoded VDJ genes. T cell populations within individuals are composed of repertoires of millions of clones each expressing a uniquely recombined configuration of TCR gene segments. Each TCR interacts with short peptide epitopes, which are derived from proteolytic turnover of cellular proteins, loaded onto major histocompatibility complex (MHC) molecules in the endoplasmic reticulum and displayed on the exterior surface. The resulting peptide/MHC (pMHC) complexes represent a sampling of the proteomic landscape of individual cells, including proteins expressed by foreign organisms, and are inspected by peripheral T cells. Individual TCR clonotypes have distinct binding preferences with respect to pMHC ligands and, if a positive TCR-pMHC interaction is formed, T cell effector functions are initiated. Collectively, the repertoire of T cells, each harboring their own TCR, provides a broad range of protection from intracellular pathogens or tumorigenesis^1^.

The semi-randomized process of TCR generation during T cell development is regulated in the thymus, where nascent TCR proteins are exposed to self-antigen pMHC in a process known as central tolerance^2^. Rearranged clonotypes responding strongly to self-antigen in the thymus are deleted, resulting in a peripheral repertoire of TCR tolerized to healthy tissue, yet poised to respond to foreign or altered peptides. In recent years, efforts to leverage TCRs as therapeutic agents^3, 4^, particularly in oncology, have been pursued by identifying TCR with demonstrated response against specific peptides from tumor associated antigens, typically from searching through patient or donor tissue, and converting these proteins into either soluble T cell engager biologics or as recombinant TCR-T cell therapies. In these therapeutic contexts, applying non-autologous, engineered, and/or supraphysiological levels of TCR circumvent natural mechanisms of central tolerance and, thus, comprehensive assessment of autoreactivity must be recapitulated in an *ex vivo* context to ensure safety and tolerability in new TCR-based therapeutics.

Profiling autoreactivity in TCR responses has remained a challenge in the field because of the tendency of these responses to be precise – capable of distinguishing between peptide sequences differing by a single amino acid^5^ – while simultaneously being highly cross-reactive, capable of responding to a vast range of millions of potential epitopes^6^. This interplay between fine discrimination and receptor-ligand promiscuity is governed by a number of biophysical and signaling phenomena such as serial triggering, kinetic proofreading, and steric interactions at the immune synapse^7–9^. Methods relying solely on affinity between TCR and pMHC, such as soluble multimeric pMHC staining reagents^10^ or pMHC yeast/viral display^11^, are often used as a proxy for functional triggering of T cell responses but run the risk of ignoring low affinity peptides with high functional potential^12, 13^. Further, animal models, even those with humanized HLA loci, are not adequate for assessing human TCR because of cross-species proteomic mismatches. Machine learning based predictive tools have recently emerged as a major area of innovation towards addressing this challenge, however, the considerable lack of functionally validated training data has meant that these approaches have not yet emerged as a practical solution^14^. Performing comprehensive functional assessment of self cross-reactivity early in the therapeutic TCR development pipeline is essential for scaling up the lead identification and pre-clinical characterization of novel TCR-based biologics and cell therapies^15^.

We have previously described a functional high-throughput screening method to profile cytotoxic T cell (CTL) responses against large libraries of short peptide-coding DNA sequences in a method we term **<u>T</u>** cell epit**<u>ope seq</u>**uencing (Tope-seq)^16^. Briefly, this method leverages a granzyme-B (GZMB)-sensitive fluorogenic reporter transgene encoded in target synthetic antigen presenting cells (sAPCs). When mixed with CTLs expressing antigen-responsive TCRs, delivery of GZMB from CTL to target sAPC – a natural response of triggered CTLs – produces a shift in fluorescence properties caused by enzymatic cleavage. The GZMB readout differs from conventional assays of T cell activation-induced markers or target cell lysis as it is a detectable early signal of T cell response spatially located within the target cell that enables precise selection of cells harboring relevant antigens out of bulk populations prior to cell loss due to apoptosis. Hence, it can be applied towards high-throughput functional screening by i) expressing large libraries of DNA-encoded minigenes, delivered as a genome integrated lentiviral vector, into reporter-expressing HLA-matched target sAPC (Fig. 1a); ii) co-culturing the library of sAPCs with CTL populations of interest; iii) sorting sAPC post co-culture by fluorescence activated cell sorting (FACS) to recover cells containing putatively antigenic minigenes (Fig. 1b); and iv) characterizing recovered cells by minigene sequencing to reveal peptide-coding sequences responsible for CTL activity (Fig. 1c). We have validated this approach in the context of randomized minigene libraries screened against primary T cells from peripheral tissue and tumor infiltrates in murine model systems^16^. We have also previously implemented the fluorogenic GZMB assay system in human cell lines and demonstrated it in the context of recombinantly expressed TCR and MHC^17^.

**Figure 1.**
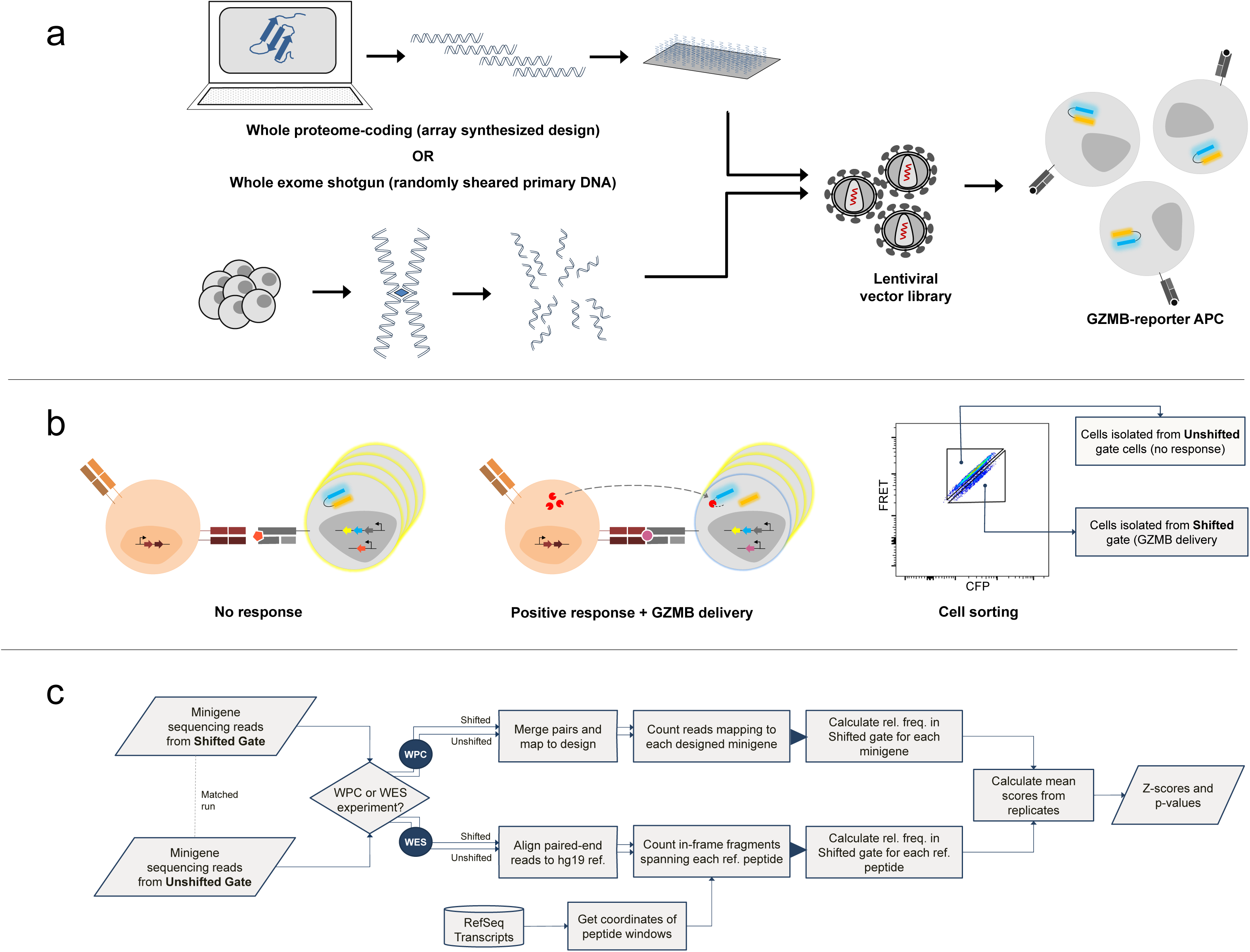
Schematic summary of Tope-seq platform. The parallel approaches for self-antigen minigene library development are illustrated in a). In one arm, the whole proteome coding (WPC) format library is created from primary amino acid sequences of all proteins identified in the human reference proteome accessed and fragmented in silico using a defined tiling scheme. The extracted amino acid sequences are then converted to DNA sequences by random codon back-translation and individually synthesized as single-stranded oligonucleotides on an array-based platform. After synthesis, DNA fragments are liberated from the array, pooled, and used to produce a lentiviral vector library. In another arm of library construction, a whole exome shotgun (WES) library was generated using primary genomic DNA from live normal human cells isolated and physically fragmented by sonication. The resultant double-stranded DNA fragments are then subjected to exome capture by RNA hybridization probes, end-polished, and adapterized for PCR amplification prior being used to produce lentiviral vector library. Library virus from either stream is used to transduce sAPC in preparation for Tope-seq based screening. The principle of the Tope-seq assay format is represented in b). Activated effector T cells transduced with the exogenous TCR-of-interest are co-cultured with sAPC harbouring an integrated self-antigen minigene library. When TCR-T cells encounter a target cell containing a minigene coding for a cognate epitope, GZMB molecules are delivered to the offending target where they cleave target-encoded ECFP-EYFP fusion proteins separated by a GZMB substrate peptide linker. The resulting loss of FRET signal (or ‘FRET-shift’) upon cleavage is monitored in flow cytometry and the library cell population is isolated by FACS to capture the Shifted cells and the counterpart Unshifted fraction. Recovered cells are characterized by targeted NGS to identify the virally encoded minigenes present in the cells from both matched gates. To quantify minigene enrichment in the Tope-seq experiment, a bioinformatic pipeline outlined in c) is used. For WPC library-based screening, matched Shifted and Unshifted gate minigene sequencing reads are paired-end merged, normalized to sample total read depth, and aligned to designed library reference sequences. Once processed, enrichment scores for each individual minigene in the library are calculated by determining the relative frequency of normalized counts in the Shifted gate (rfs) as a proportion of the total across both gates. For WES libraries, processing is performed by mapping paired-end reads to the reference human genome and using a sliding reference peptide window to scan all k-mer size ranges (from 8 to 11 residues in length) across the coding genome. In-frame minigenes traversing each window are counted and normalized to read depth for each matched gate dataset prior to calculating rfs score for individual peptides, rather than producing minigene-specific scores. Where applicable, rfs scores from replicate experiments are calculated as the geometric mean of all measurements. We defined two separate read processing pipelines for the two formats of minigene libraries since WES libraries contain epitopes embedded within an undefined number of possible distinct minigene fragments, WES fragments are generally longer in length than would allow for an accurate paired-end merging step, randomly sheared WES fragments could be cloned into viral vector in one of six possible expression frames (whereas frame is precisely controlled in WPC format), and WPC minigenes are constructed with non-native codon usage.

In this study, our goals were to apply Tope-seq technology to the characterization of TCR autoreactivity, while addressing the challenge of background signal arising from endogenous antigens. Here, we apply the platform towards comprehensive functional characterization of a3a (an *ex vivo* affinity-enhanced TCR that previously failed in clinical trials due to cardiotoxicity mediated by unexpected cross-reactivity^18^) and its wild-type thymically-selected counterpart (EB81.103^19^) against large libraries of potential epitopes representing all possible genome encoded peptides. We show that the therapeutic target epitope, EVDPIGHLY, derived from the MAGEA3 protein was detectable for both TCRs by Tope-seq and the cross-reactive epitope previously determined to be the cause of clinical failure, ESDPIVAQY, derived from the Titin (TTN) protein, was detected in analysis of the affinity-enhanced a3a only. We also report strategies for developing libraries of potential epitopes, contending with background from endogenously expressed proteins, and performing bioinformatic analyses on large Tope-seq datasets.

## Results

### Development of minigene libraries spanning self-antigen space

To conduct Tope-seq screening on complete self-antigen minigene libraries, we first needed to develop suitable libraries to express in reporter target cells. One approach employed to this end was a whole proteome coding (WPC) minigene synthesis strategy, accomplished by designing *in silico* a set of amino acid sequences representing all possible peptides potentially expressed by the human coding genome. To do this, each protein coding sequence in the human reference proteome was deconstructed *in silico* by extracting tiled amino acid segments and converting these to DNA minigenes by backtranslation. Redundant protein domains observed in protein isoforms or across highly homologous proteins were consolidated prior to tiling to avoid overrepresentation of peptide subsequences in the final fragment library. The resulting set of >5 x 10^5^ minigenes was array synthesized (Twist Biosciences) to yield a single-stranded DNA oligonucleotide pool, which we then cloned into lentiviral transfer plasmid. The prepared WPC library plasmid was characterized by NGS analysis of minigenes amplified directly from starting library plasmid stock, which showed >97% of reads successfully mapped to the designed reference minigenes and >97% of the designed library was represented, indicating a very low incidence of introduced artifacts or sequence dropout (Suppl. Tab. 1).

We then proceeded to use prepared plasmids to generate lentiviral vector and transduce K562 cells previously engineered to express a HLA-A*01:01/GZMB-FRET reporter gene cassette (referred to as KFRET.A0101 cell line)^17^. Our initial transduction strategy was a low MOI (MOI=0.2) approach to favor probabilistically the production of no more than a single minigene integration per cell. After transduction and purity sorting, we assessed the integration of WPC minigene libraries by performing NGS analysis of minigene amplicons recovered from cells (Suppl. Tab. 2). On average, replicate samples were found to contain 86.9% of designed minigene sequences. Interestingly, however, when considering the union of all four replicates, the increased depth from combining the data resulted in library coverage roughly similar to the source plasmid. We reasoned, therefore, that bottlenecking was not introduced at the transduction step since sampling more cells resulted in near complete coverage.

Based on this finding, we repeated WPC minigene library transduction to increase the representation of minigenes in the cell-expressed library. A second iteration of KFRET.A0101.WPC libraries were produced by increasing library transduction MOI (MOI = 7) and permitting multiple minigene copies per target cell. Although there is risk in a high MOI approach of passenger minigenes becoming captured in the FRET-shifted minigene population and falsely detected in Tope-seq, we expected this effect would be buffered by conducting transduction at a sufficiently large scale to ensure clonal redundancy of each minigene and render it unlikely that any true negative passenger minigenes would repeatedly co-occur with true positive epitopes in the founder clone population. We assessed the integration of high MOI WPC minigene libraries post-transduction (Suppl. Tab. 3) and determined that each high MOI replicate produced represented >97% of the designed minigene library. Thus, the high MOI library strategy was carried forward for subsequent Tope-seq experiments.

In parallel to synthetic WPC library development, we undertook an alternative strategy towards the generation of minigene libraries for expression in target reporter cells. We explored primary nucleic acids as source material from which minigene libraries can be built, in this case constructing a comprehensive self-antigen minigene library by using normal human genomic DNA from a mixture of multiple primary healthy donors. The genomic starting material was acoustically sheared and subject to exome capture and adapterization. These steps were performed off-site (Genewiz) using standardized protocols for Illumina library construction and, rather than proceeding with exome sequencing, we interposed minigene vector library cloning and target cell line creation to prepare a Tope-seq-ready exome minigene library. Since, for this format, we did not have a pre-defined expected library size as was the case with the WPC minigene library, we forewent QC sequencing at the plasmid level and, instead, assessed library diversity by sampling transduced cells and performing NGS analysis of minigene amplicons recovered therein (Suppl. Tab. 4). The results of this analysis indicated that the starting whole exome shotgun (WES) cell library represented >2.5 x 10^7^ unique minigene fragments (or greater than 100X coverage of the human exome). Notably, the diversity of WES format libraries was considerably larger than WPC libraries. This was expected since WES fragments were generated by shearing DNA at random breakpoints and inserted into lentivector cassette in six possible reading frames.

### Initial library screen in sAPC with endogenous epitope was unsuccessful

In previous work, we observed FRET-shift signal during co-culture of both EB81.103 and a3a TCR-expressing T cells against KFRET.A0101 cells in the absence of any introduced minigene^17^. This suggested the presence of endogenous MAGEA3 protein and potentially other self-antigens in these cells, which we suspected could represent a source of background for library-based screening experiments. Epitopes derived from host cell proteins would render all cells in the target population selectable by FRET-shift FACS, irrespective of the transduced minigenes contained within. We sought to assess to what extent this background would confound the interrogation of exogenous minigene libraries and, by extension, whether the known reactivities of the test TCRs could still be detected despite the presence of endogenous antigen.

Initial screens were performed by co-culturing a3a CD8^+^ TCR-T cells with KFRET.A0101 cells containing either WPC or WES minigene library. For each screening experiment, FRET-Shifted and Unshifted target cell fractions were isolated according to the gating strategy illustrated in Suppl. Fig 1. Immediately after cell capture, cells recovered from each gate were separately processed for genomic DNA purification. Minigenes integrated in captured genomic DNA samples were recovered by targeted PCR and sequenced. After minigene sequencing, data was processed via Tope-seq analysis pipelines developed specifically for either WPC or WES experiments (Fig. 1c). Enrichment was measured by computing the **<u>r</u>**elative **f**requency of each individual library sequence’s normalized count in the **<u>S</u>**hifted gate sample as a proportion of its total normalized count across both gates (rfS). Each rfS score, a sampling proportion of the library population, is normally distributed (Suppl. Fig. 2). Hence, Z-scores and p-values were produced for each minigene or reference peptide sequence for significance testing.

Analysis of results from WPC experiments conducted in duplicate revealed that minigenes containing the known MAGEA3 and TTN derived epitopes were not significantly enriched above background. Similarly, our WES experiments also indicated that the positive control epitopes could not be detected from background under these conditions (Fig. 2). This provided evidence that T cell response to non-library encoded epitopes was outpacing response to epitopes derived from our exogenous minigene libraries. To investigate if this was the case, and also rule out limit of detection as the explanation for this result, we repeated library screens in sAPC cell chassis lacking endogenous epitopes derived from host cell proteins.

**Figure 2.**
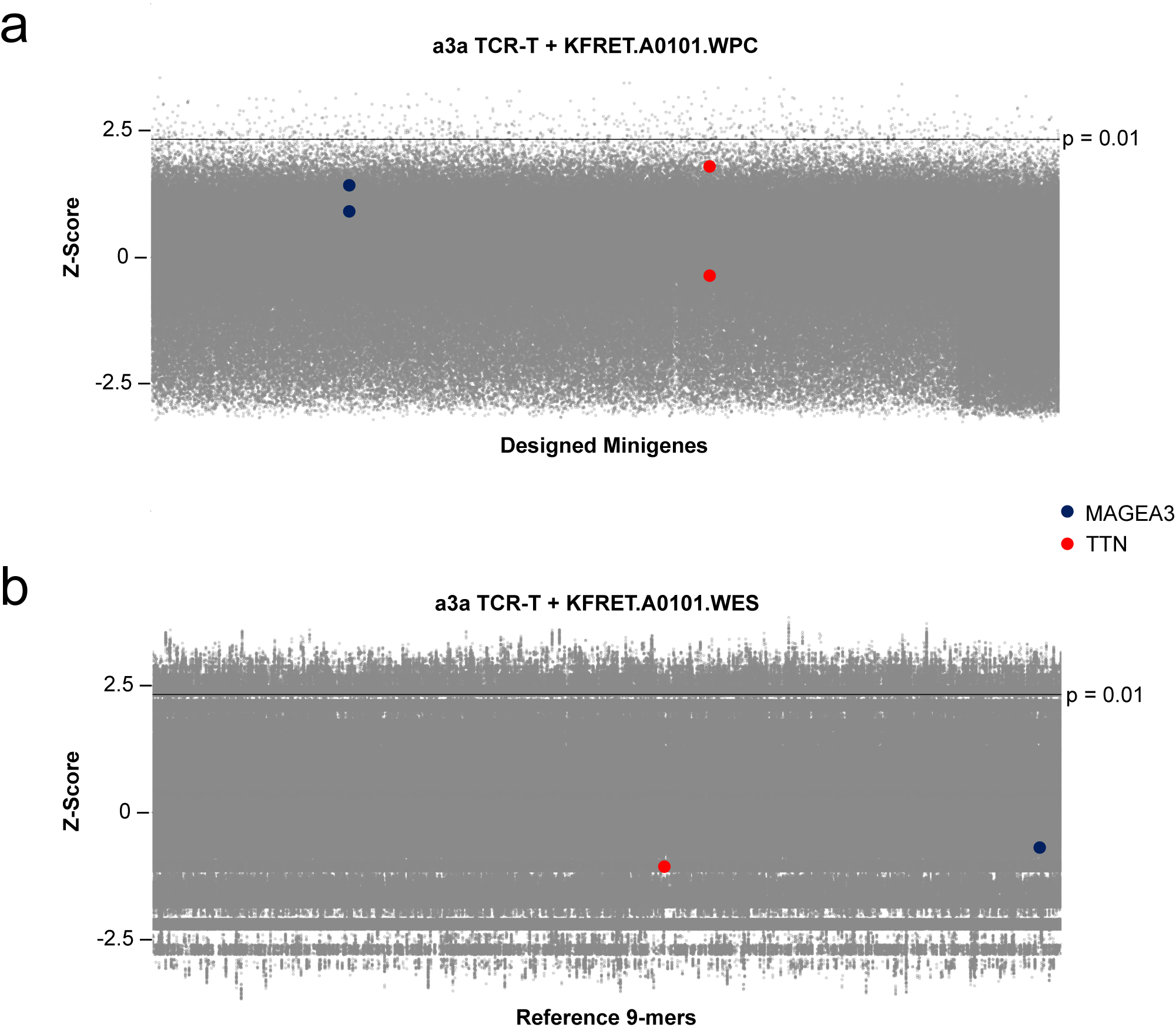
Tope-seq assessment of a3a TCR-T using unedited K562 target chassis. Tope-seq readout from initial test of a3a TCR-T cells co-cultured with KFRET.A0101 cells containing either the WPC library (a) or WES library (b). In both experiments, an effector/target ratio of 1:1 was used to assemble co-cultures. For each screen, >6 x 10^7^ target library cells were incubated with TCR-T cells for a period of 12 hours prior to FACS isolation of Shifted and Unshifted cells. Recovered cells were subject to gDNA purification and minigene amplification before Illumina sequencing. The frequency of reads in Shifted relative the normalized read counts in Shifted + Unshifted for each minigene (for (a)) or 9-mer coding genomic locus (for (b)) were computed and used to calculate Z-scores plotted on y-axes. Each grey point represents an individual minigene or 9-mer peptide coding locus. The control MAGEA3 and TTN epitope-containing minigenes (for a) or minimal epitopes from MAGEA3 and TTN (for b) are highlighted according to the figure legend. Significance threshold representing p = 0.01 is indicated on y-axes as a solid line (Z-scores above line are p < 0.01).

### Alternative target cell chassis circumvent endogenous background

Upon observing that the monitored test antigens could not be distinguished above background using target cell lines harboring native reactivities of the EB81.103 and a3a TCRs using either of our minigene library formats, we theorized that strategies to avoid background from endogenous antigen would enable detection of positive control epitopes from complex self-antigen libraries. In previous work, we developed a CRISPR/Cas9 edited version of the KFRET.A0101 cell line, in which the EVDPIGHLY epitope within the *MAGEA3* gene at the native locus was deleted^17^. We found that this genetic intervention ablated background FRET-shift response in co-culture with EB81.103 TCR-T cells. However, despite genome-level removal of the target epitope in the base cell line, there was no difference in signal in flow cytometry between co-cultures of a3a TCR-T cells and KFRET.A0101.MAGEA3*^-/-^* with or without re-introduced MAGEA3 minigene. We reasoned that this likely was due to the presence of other potential cross-reactive epitopes of a3a endogenously expressed in the K562 cell line.

To confirm this hypothesis, we validated, in this study, an alternative “clean” target cell chassis to use as a host cell line for minigene library screening by Tope-seq. To this end, we generated sAPC based on 721.221, which is an EBV-transformed B-lymphoblastoid cell line rendered MHC-null by gamma-irradiation mutagenesis^20^. Similarly to K562, despite loss of its own MHC class I α-chain expression, 721.221 cells retain *B2M* expression and the necessary antigen-processing and presentation machinery to generate surface pMHC complexes when provided with

HLA-I coding sequence. We converted base 721.221 cells into 721FRET.A0101 cells by viral transduction to equip them with HLA-A*01:01 coding sequence and the FRET-reporter transgene. We subsequently co-cultured these targets, either without minigene or further transduced with MAGEA3 minigene, with a3a TCR-T cells to determine FRET-shift signal-to-background level. The results of this testing (Fig. 3, Suppl. Fig. 3) indicated 721FRET.A0101.MAGEA3^143–202^ cells resulted in minigene-specific response distinguishable in FRET-shift assay from the negative control, demonstrating that expressed self-epitopes in K562 that confounded assessment of the a3a TCR are not also expressed in the base 721.221 cell line.

**Figure 3.**
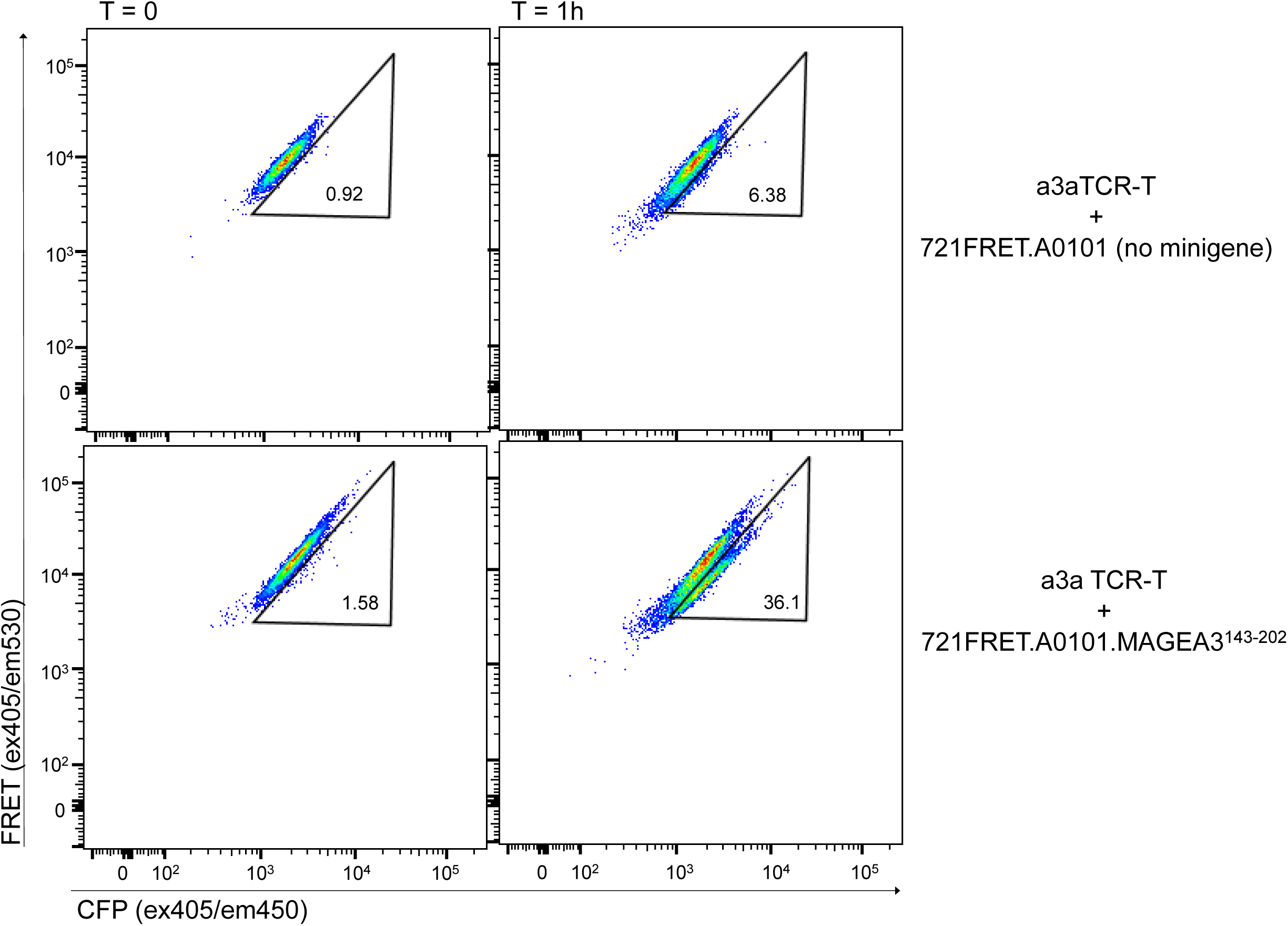
721.221 based target cells do not trigger endogenous background response from a3a. 721FRET.A0101 and 721FRET.A0101.MAGEA3^143–202^ target cells were co-cultured with primary cytotoxic T cells freshly isolated by FACS purification CD8^+^ CD4^-^ CD56^-^ T cells from healthy donor PBMC, activated by plate-bound anti-CD3/28 stimulation, and transduced with a3a TCR lentivirus. The resultant a3a TCR-T cells were co-cultured with targets at a 1.5:1 ratio (to compensate for 68% TCR transduction efficiency) for 1 hour and analyzed by flow cytometry to quantify GZMB-induced FRET-shifting in targets with or without MAGEA3 minigene.

### Comprehensive self-peptide assessment using WPC library

Based on these results, we repeated Tope-seq analysis of a3a TCR-T cells against WPC minigene libraries expressed in the 721FRET.A0101 target cell chassis. In parallel, we also set out to investigate wild-type EB81.103 TCR-T cells against WPC library encoded in KFRET.A0101.MAGEA3*^-/-^* cells (the native *MAGEA3* gene knockout line). To prepare for these experiments, new target library populations were produced in their respective cell chassis by performing lentiviral transduction at the same MOI and total scale as the previous WPC experiment. Upon library cell line purity-sorting and TCR-T production, Tope-seq experiments were conducted in duplicate for each condition. FRET-shift FACS sorting and minigene sequencing was performed identically to initial screening experiment.

The results of these Tope-seq screens indicated new cell libraries represented >97% of library sequences, similar to previous versions, and that positive control minigenes could be successfully detected in both experimental conditions. In the EB81.103 TCR screen, both MAGEA3 minigenes containing the EVDPIGHLY epitope were significantly enriched above background (p < 0.001, rank >99.99^th^ percentile) while both TTN minigenes containing the ESDPIVAQY epitope were not significantly enriched (Fig. 4a). In the a3a TCR screen, the MAGEA3 minigenes were significantly enriched (p < 0.001, >99.8^th^ percentile) while, in this case, both TTN minigenes were also found to be significantly enriched (p < 0.001, rank 99.8^th^ percentile) (Fig 4b). These results reflect the findings made in the original clinical case report after severe off-target cross-reactivity was observed *in vivo and* show that comprehensive self-antigen screening of a whole-proteome coding library with Tope-seq would have raised this cross-reactivity as a risk factor during the discovery and development phases of the a3a TCR. This result also confirmed our hypothesis that eliminating response to host cell antigens is sufficient for enabling Tope-seq based assessment of TCRs-of-interest against complex self-antigen libraries.

**Figure 4.**
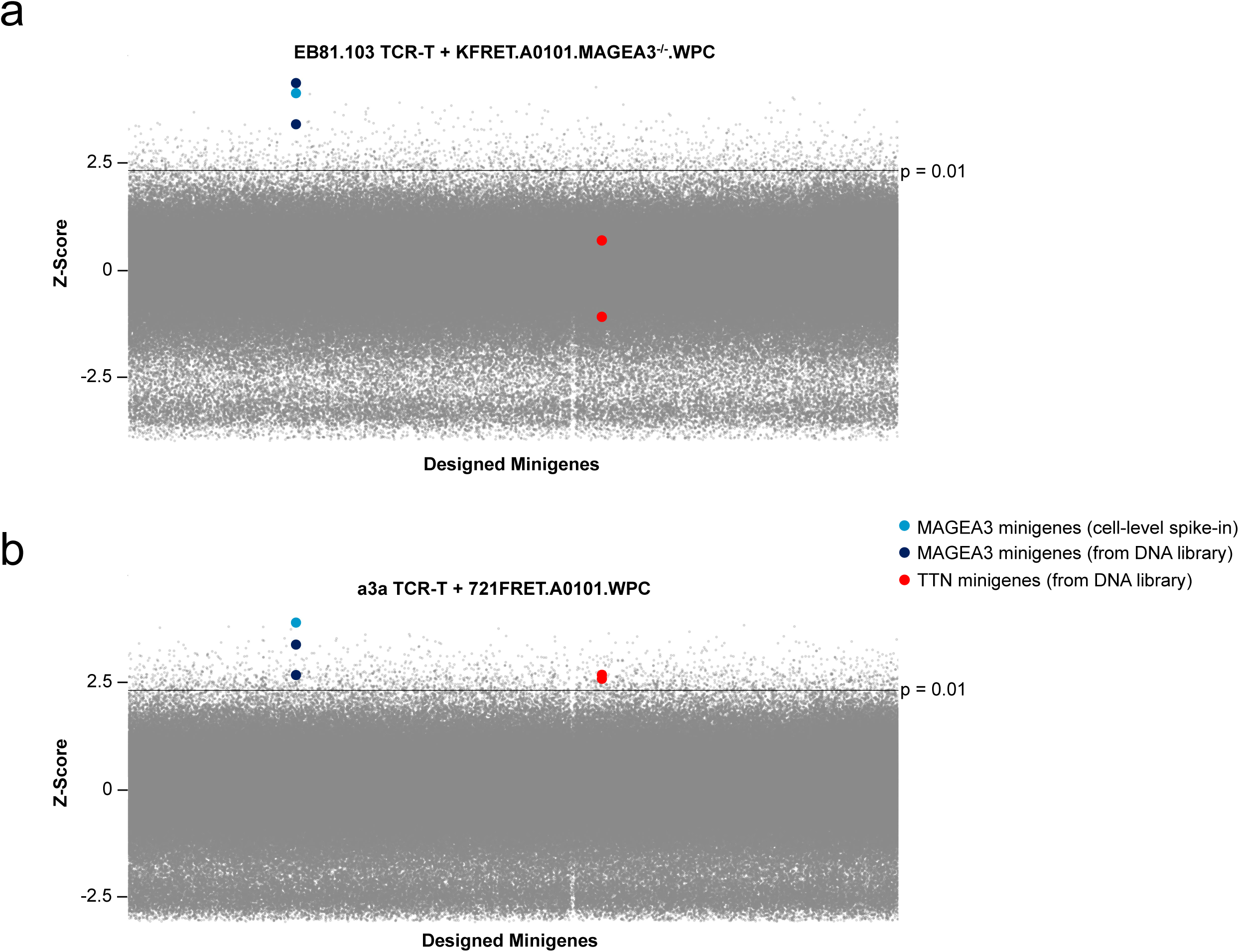
Tope-seq assessment of EB81.103 and a3a TCR against WPC library in clean target chassis. Tope-seq readout from initial test of EB81.103 TCR-T cells co-cultured with CRISPR-edited KFRET.A0101.MAGEA3^-/-^ cells containing WPC library (a) or a3a TCR-T cells co-cultured with 721FRET.A0101.MAGEA3^-/-^ cells containing WPC library (b). In both experiments, an effector/target ratio of 1:1 was used to assemble co-cultures. For each screen, >6 x 10^7^ target library cells (spiked with 1.5 x 10^3^ KFRET.A0101.MAGEA3^-/-^.MAGEA3^143–202^ re-introduced minigene cells as an internal control) were incubated with TCR-T cells for either a period of 12 hours (for KFRET cells) or 1 hour (for 721FRET cells) prior to FACS isolation of Shifted and Unshifted cells. Recovered cells were subject to gDNA purification and minigene amplification before Illumina sequencing. Both experiments were conducted in duplicate. The frequency of reads in Shifted relative the normalized read counts in Shifted + Unshifted for each minigene (for panel a) or 9-mer coding genomic locus (for panel b) were computed in each experiment. Geometric mean of minigene scores across both replicates were then calculated and used to calculate Z-scores plotted on y-axes. The control MAGEA3 and TTN epitope-containing minigenes are highlighted according to the figure legend. Significance threshold representing p = 0.01 is indicated on y-axes as a solid line (Z-scores above line are p < 0.01).

### Bioinformatic refinement of Tope-seq data by clustering

We applied further bioinformatic analysis to investigate other minigenes enriched above the p < 0.01 threshold in WPC experiments (n = 1,850 in EB81.103 experiment; n = 2,283 in a3a experiment). Interestingly, in the original unsuccessful a3a TCR-T/WPC library experiment in unedited sAPC, 3-fold fewer (n = 758) minigenes were found to meet the statistical threshold. Our goal was to examine whether putative cross-reactivities arise from common epitope motifs or if multiple epitope clusters emerged for either of the TCRs interrogated. To do this, we analyzed all 8- to 11-mer peptide sequences embedded in each minigene for both experiments using a clustering approach. As a first step, initial filtering of peptide lists was performed by removing peptides from non-significant (p > 0.01) minigenes. We then determined, for peptides with occurrences in multiple different minigenes, which sequences met the statistical significance threshold more frequently than would be expected due to chance (binomial exact test). After initial filtering, peptide sequences were further filtered by performing peptide-MHC binding prediction using NetMHC 4.0^21^. To avoid introducing undue bias in the analysis, we applied a very low stringency cutoff (rank score < 5). Peptides were then grouped using GibbsCluster 2.0^22^, an algorithm for simultaneous alignment and clustering based on a Gibbs sampling approach for deconvoluting motifs in complex sets of sequences. Motif detection was performed by producing a baseline distribution of motifs from randomly generated peptides and conducting significance testing of GibbsCluster scores from each experiment-derived motif against the distribution of scores from baseline random peptide motifs (Fig. 5a).

**Figure 5.**
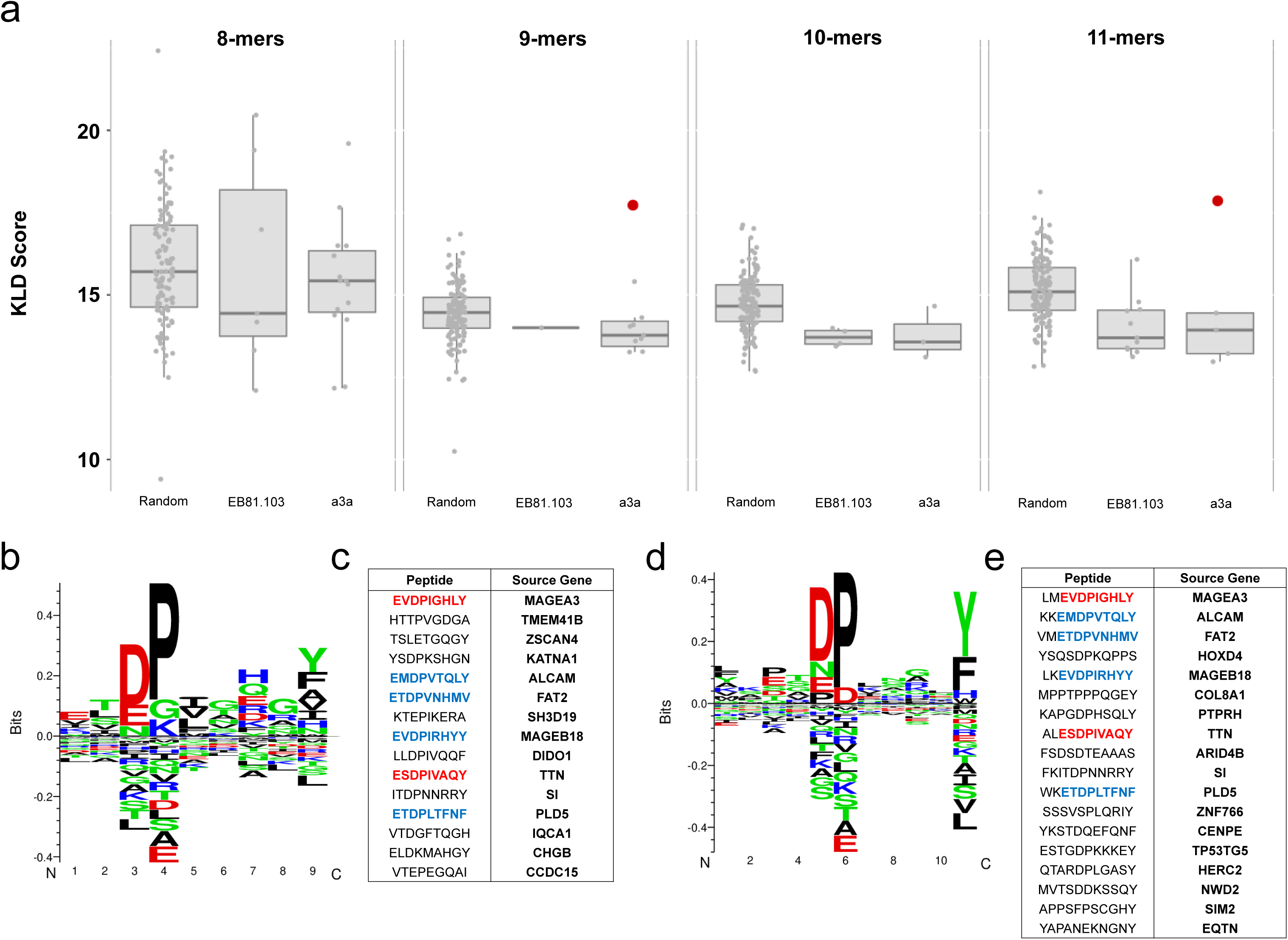
Clusters of similarity emerge among hits from a3a TCR-T WPC screening. a) Minigenes in EB81.103 TCR-T + KFRET.A0101.MAGEA3^-/-^ and a3a TCR-T + 721FRET.A0101 WPC library experiments were broken into k-mers of sizes ranging from 8-11 amino acids and filtered on Z-score threshold and NetMHC 4.0 binding (using a relaxed threshold of rank < 5). Filtered peptide lists were used as input sequences for GibbsCluster 2.0, testing at various K-means clustering ranging from 1-15 clusters. For each experiment of each peptide size, the K-means regime producing the highest average Kullback-Leibler Divergence (KLD) score was carried forward for further analysis (individual clusters plotted here with interquartile boxplot overlay). For significance testing, clusters from each solution were compared to a baseline distribution generated from clustering randomized peptides (all filtered to meet rank score < 5 in NetMHC 4.0). The random baseline represents cluster scores from the top performing solution in ten iterations of random peptide clustering and was found to be approximately normally distributed by Kolmogorov-Smirnov test. Only two clusters (highlighted in red) were found to be statistically significant compared random peptide background (p < 0.01 by one-sample Z-test). These clusters were found in the 9-mer and 11-mer analysis of the a3a TCR-T experiment. Motifs of the significant clusters are shown in b) and d) and the core alignment peptides for each cluster are shown in c) and d), respectively. Amino acids highlighted in red represent known test peptides from MAGEA3 and TTN. Amino acids highlighted in blue represent peptides previously observed in other analyses of the a3a TCR^23^. Non-highlighted represent peptide hits not previously observed in orthogonal studies. Peptides are shown listed with source human genes.

Overall, only two motifs emerged with statistically significant GibbsCluster scores, both arising from the a3a TCR dataset (Fig. 5b, d). This was not unexpected given that a3a is a known promiscuous and self-reactive engineered TCR while EB81.103 is a thymically-selected TCR that would not be expected to form clusters of reactivity in a self-antigen library. We also observed that the two emergent motifs were derived from different peptide size analyses but contained notable overlap in the core peptides used to generate them. The discovered motifs were observed to contain the known positive epitopes from MAGEA3 and TTN, as well as additional reactivities that had been previously reported in prior studies^23^. We also detected additional reactivities that were not previously reported and may represent newly discovered self-epitopes of the a3a TCR (Fig. 5c, e), some of which have high tissue-specific expression in Human Protein Atlas^24^ reference data (Table 1) and may also potentially be clinically relevant. Some putative WPC hits from a3a screening were also shown to be expressed in Human Protein Atlas data for the K562 cell line. These potential K562-specific antigens are expressed at relatively lower levels in 721.221 cells^25^, providing evidence supporting the hypothesis that some of the detected putative epitopes from this experiment were contributing to background signal in K562 chassis-based library screens.

**Table 1.**
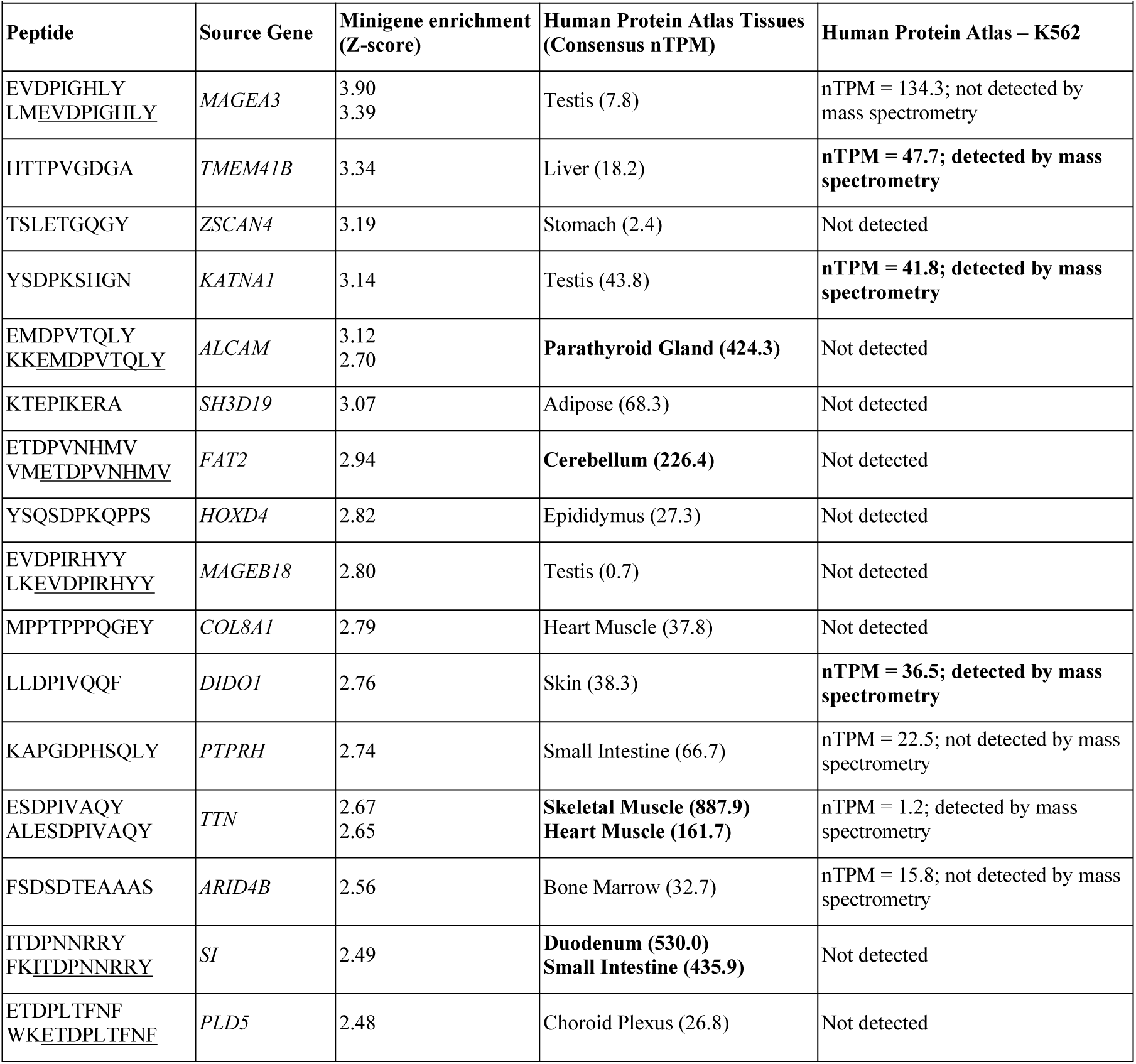
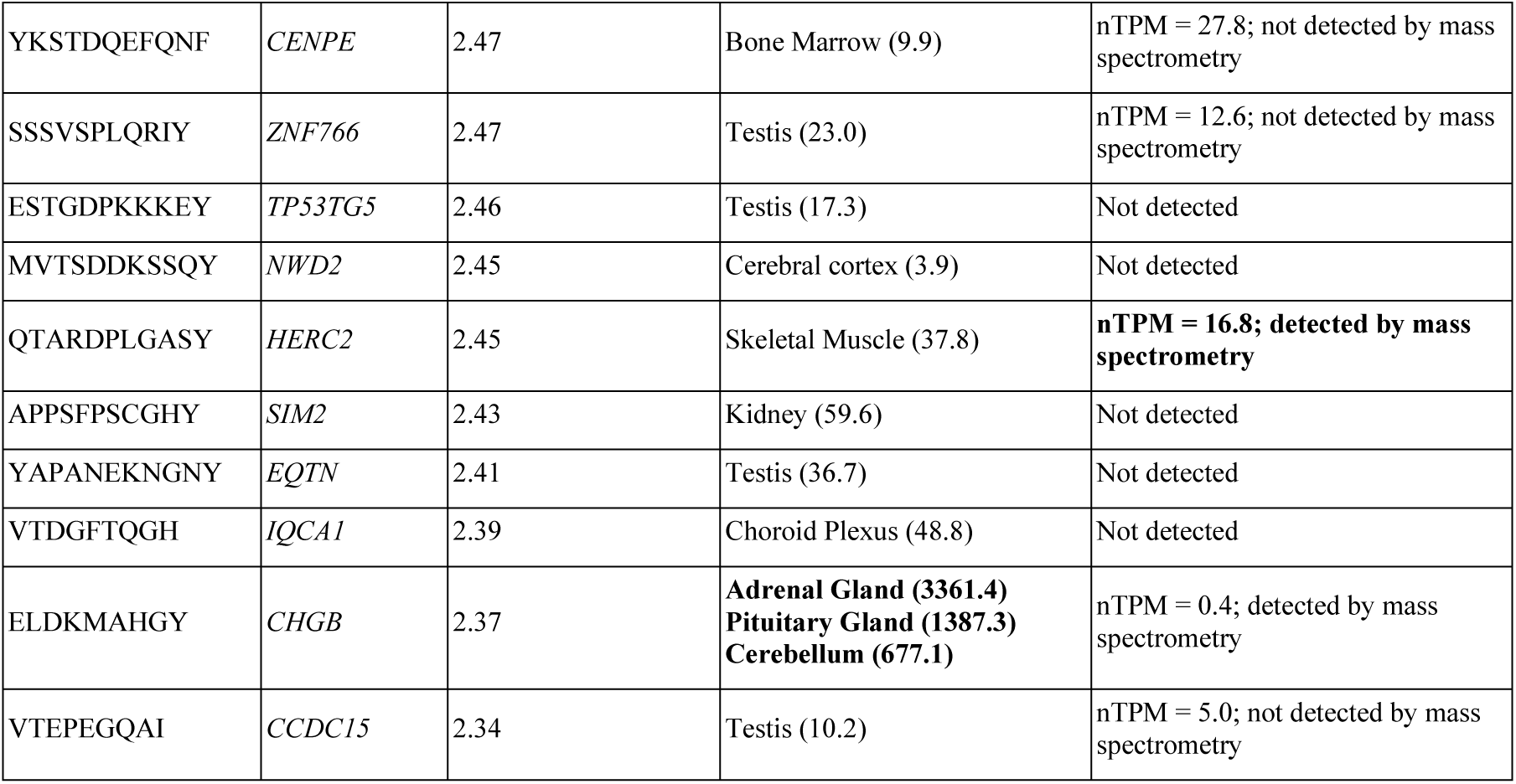
Tissue- and K562 cell line-specific expression of putative self-antigen hits from a3a TCR-T WPC screen. 9-mer and 11-mer peptides, comprising the core alignments of detected epitope motifs derived from Tope-seq analysis of a3a TCR-T, were cross-referenced with respect to their source antigenic protein to the Human Protein Atlas dataset. Peptides with complete overlap were grouped together for analysis. The source genes of each putatively immunogenic epitope and Z-scores from library minigenes containing epitopes are shown. For each, the consensus normalized transcript per million (nTPM) values for the tissue type with the highest reported transcript level were retrieved from the Human Protein Atlas and shown here. Multiple tissue types were reported (in bold) for cases where transcript level was higher than expression level of *TTN* in heart muscle, which was determined to be clinically relevant in the original case report. Expression level of motif core alignment peptides was also investigated for K562 data included in the Human Protein Atlas cell line repository. The nTPM values, where applicable, are shown below as well as presence or absence scoring at the protein level as determined by mass spectrometry.

### Biopanning as a strategy for overcoming endogenous reactivities

Our strategies for deleting endogenous epitopes by genome editing or developing alternative clean target cell chassis to prepare a suitable library host for Tope-seq based self-antigen assessment were effective in this study. However, we considered that these options may not always be available, particularly in cases where putative self cross-reactivities are unknown or are in essential genes that cannot be knocked out or otherwise avoided. Therefore, in addition to exploring engineered or alternative target cell lines for avoiding background from endogenous antigens, we sought to investigate whether this background could be overcome by iteratively re-screening selected minigenes to enrich epitopes progressively, analogous to biopanning techniques used in phage-display technologies^26^. Because of the large diversity of unique minigenes in WES libraries, as compared to WPC, we performed testing of biopanning in these libraries anticipating that significant bottlenecking of the library diversity between first and second round experiments would enable us to see enrichment effects clearly. Our hypotheses were twofold. We first expected that re-screening leftover Shifted gate amplicon produced during NGS library preparation from a 1^st^ round experiment would result in increasing enrichment scores of known positive epitopes. We also expected that most peptides with significant enrichment scores from the first round would be undetected or non-significant in the second round.

We constructed 2^nd^ round WES libraries by re-adapterizing Shifted gate minigenes from 1^st^ round screens of both TCRs for cloning by PCR and inserted them into library expression lentiviral transfer plasmid. Cloned 2^nd^ round libraries were used to generate lentiviral vector and create library cell lines in non-CRISPR edited KFRET.A0101 cell chassis. Sampling the prepared 2^nd^ round libraries for NGS QC revealed that both libraries contained 1-2 × 10^6^ unique minigenes (representing 4-8% of the minigenes contained in the original library) (Suppl. Tab 5). Tope-seq experiments were conducted for TCR-T cell preparations containing either EB81.103 or the a3a TCR against their respective 2^nd^ round library. Our results indicated that, in the EB81.103 TCR experiment, the EVDPIGHLY epitope from MAGEA3 was significantly enriched above background (p < 0.001, rank >99.99^th^ percentile) after 2 rounds while the ESDPIVAQY epitope from TTN was not significantly enriched (Fig. 6a). In the a3a TCR screen, the MAGEA3 epitope was significantly enriched (p < 0.001, rank >99.9^th^ percentile) while the TTN epitope was also found to be significantly enriched (p < 0.01, ranked >95^th^ percentile) after 2 rounds (Fig. 6b). These results were similar to what we observed in the analysis of WPC library in clean background cells, indicating that iterative re-screening, or biopanning, can also be deployed as a tool, either independently or potentially in conjunction with chassis editing strategies, for contending with background arising from naturally occurring antigens in live host sAPC.

**Figure 6.**
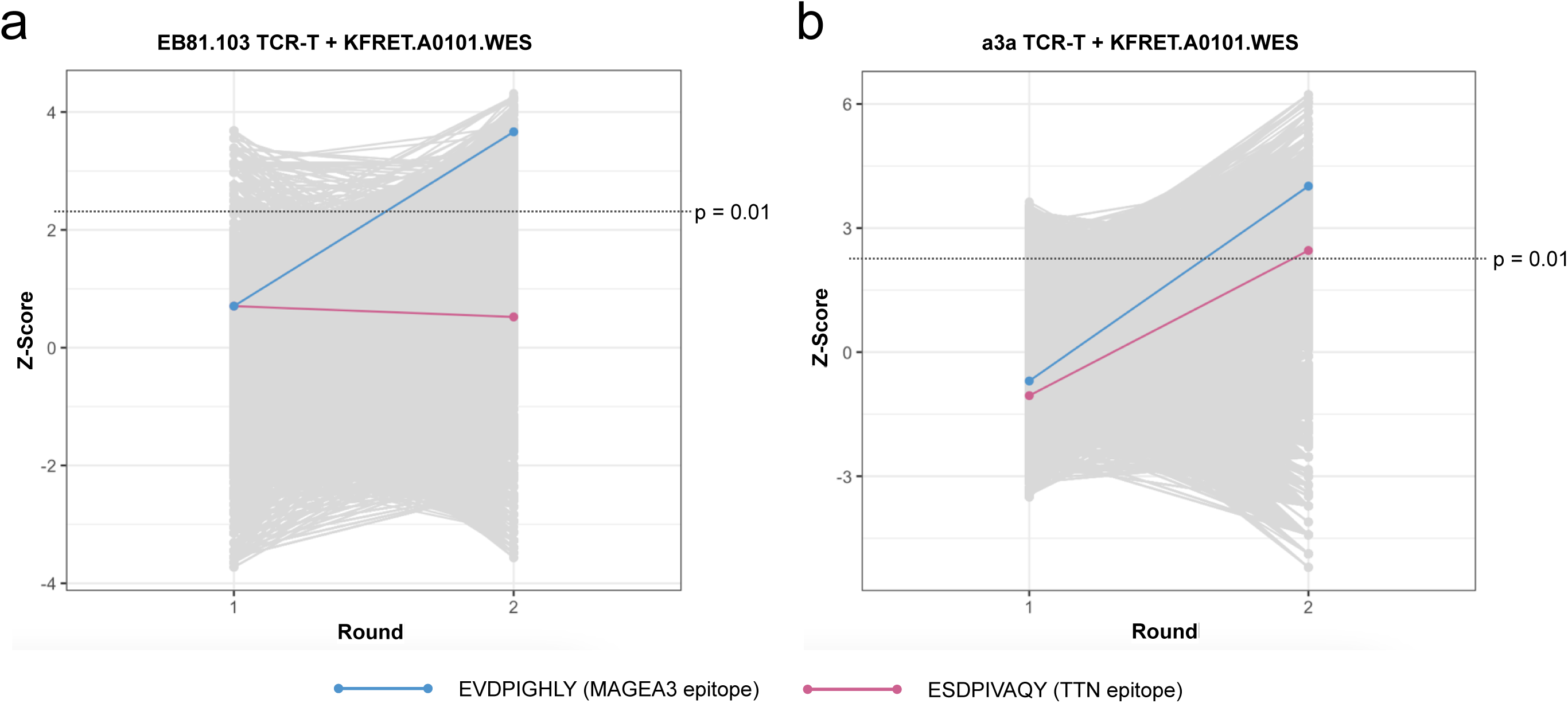
Enrichment observed in 2^nd^ round panning screen of WES library. Tope-seq readouts from 1^st^ and 2^nd^ round screens of a) EB81.103 TCR-T cells and b) a3a TCR-T cells against non-CRISPR edited KFRET.A0101 cells containing WES library (for 1^st^ round experiments) or respective re-cloned panning library (for 2^nd^ round experiments). In both experiments, an effector/target ratio of 1:1 was used to assemble co-cultures. For each screen, >6 x 10^7^ target library cells were incubated with TCR-T cells for a period of 12 hours prior to FACS isolation of Shifted and Unshifted cells. Recovered cells were subject to gDNA purification and minigene amplification before Illumina sequencing. The frequency of in-frame reads spanning each 9-mer coding genomic locus in Shifted relative the normalized read counts in Shifted + Unshifted were computed and used to calculate Z-scores plotted on y-axes. Each peptide scored in the 1^st^ and 2^nd^ rounds are plotted as a grey point in each group and connected between rounds by a line to indicate increase or decrease after panning. The control minimal epitopes from MAGEA3 and TTN are highlighted according to the figure legend. Significance threshold representing p = 0.01 is indicated on y-axes as a solid line (Z-scores above line are p < 0.01).

We next sought to investigate broad patterns among other enriched peptides in the dataset. To this end, we scored peptides from 8-mer, 10-mer and 11-mer reference window settings. Strikingly, we noted that for all analyses, there were >10-fold more peptides meeting significance threshold after both rounds in the a3a experiment than in EB81.103 (Table 2), perhaps reflecting the highly promiscuous nature of the a3a TCR relative to its wild-type, thymically-selected counterpart. Repeating the filtering and clustering analysis described above for the WPC experiments, we were unable to derive new statistically significant peptide motifs from the WES library format. Since the 2^nd^ round WES panning libraries still represented a substantially larger number of unique minigenes than are present in the WPC library, we considered that additional panning would be necessary to detect clusters of reactivity by this method. We also observed that most peptides (>93% for all analyses) meeting the p-value cutoff in the 1^st^ round were not re-discovered in the 2^nd^ round, supporting our hypothesis that the panning approach is useful for removing background. However, a larger proportion of 1^st^ round hits were re-discoverable in the 2^nd^ round than would be expected due to chance (1.4% and 6.3% for EB81.103 and a3a 9-mer analyses, respectively; p < 0.001 by one-tailed exact binomial test). Additionally, although the dataset was approximately evenly split between peptides increasing in score (incliners) and those decreasing between rounds (decliners), a significant proportion of the incliner population was found to meet the cutoff in 2^nd^ round screening for both experiments (1.6% and 13.3% for EB81.103 and a3a 9-mer analyses, respectively; p < 0.001 by one-tailed exact binomial test), which provided evidence that positive enrichment of immunogenic epitopes was occurring.

**Table 2.** Summary of putative enrichment dynamics in two rounds of Tope-seq biopanning of WES self-antigen library. Experiment results from both the EB81.103 (wild-type) and a3a (engineered) TCR variants were processed for all peptide sizes ranging from 8-11 amino acids in length. Shown, for each experiment and size analysis, are the number of peptides detected in each round (as well as the number of peptides meeting significance threshold after two rounds), the percentages of peptides that increased or decreased in rfS score between rounds, the percentages of significant peptides from the 1^st^ round also found to be significant in the 2^nd^ round, the percentage of significant peptides from round 2 that increased in rfS score between rounds, and the percentage of non-significant peptides from the 1^st^ round that became significant after the 2^nd^ round.

|  |  |  |  |  |  |  |  |  |
| --- | --- | --- | --- | --- | --- | --- | --- | --- |
|  | **EB81.103** |  |  |  | **a3a** |  |  |  |
|  | 8mers | 9mers | 10mers | 11mers | 8mers | 9mers | 10mers | 11mers |
| # Peptides 1<sup>st</sup> rd. | 5,299,213 | 5,161,203 | 5,023,895 | 4,887,440 | 5,140,672 | 5,005,841 | 4,871,757 | 4,738,480 |
| # Peptides 2<sup>nd</sup> rd. | 1,673,140 | 1,620,446 | 1,568,225 | 1,516,499 | 1,739,540 | 1,682,888 | 1,626,736 | 1,571,267 |
| % Peptides with 1<sup>st</sup> rd. score < 2<sup>nd</sup> rd. score (incliners) | 41.9 | 42.0 | 42.1 | 42.1 | 50.6 | 50.8 | 50.9 | 51.1 |
| % Peptides with 1<sup>st</sup> rd. score > 2<sup>nd</sup> rd. score (decliners) | 58.1 | 58.0 | 57.9 | 57.9 | 49.4 | 49.2 | 49.1 | 49.9 |
| % Sig. peptides ($p < 0.01$) from 1<sup>st</sup> round re-discovered in 2<sup>nd</sup> | 1.8 | 1.4 | 1.3 | 1.2 | 7.3 | 6.3 | 5.5 | 5.0 |
| # Sig. peptides ($p < 0.01$) 2<sup>nd</sup> rd. | 10,823 | 10,635 | 10,437 | 10,283 | 124,458 | 120,367 | 116,284 | 112,250 |
| Incliners as % of 2<sup>nd</sup> rd. sig. peptides | 99.4 | 99.3 | 99.3 | 99.2 | 94.5 | 94.5 | 94.4 | 94.4 |
| % Incliners n.s. in 1<sup>st</sup> rd. & sig. in 2<sup>nd</sup> rd. | 1.53 | 1.55 | 1.57 | 1.60 | 13.3 | 13.3 | 13.3 | 13.2 |

## Discussion

A key lesson taken from the original case report of the catastrophic failure of the a3a TCR^27^ is the urgent need for functional assessment of engineered therapeutic TCR against the widest panel of possible self-antigens to rule out dangerous off-target cross-reactivities early in therapeutic development. In this study, we used the a3a TCR as a test case to evaluate the utility of Tope-seq in directly screening all peptides encoded in the genome for their potential to elicit autoreactivity.

Currently. the most common *in vitro* functional assay for probing TCR cross-reactivity is based on peptide-pulsing target cells with pools of partially randomized peptide to deduce putatively important residues necessary for TCR triggering^23, 28–30^. The main drawbacks of this technique are that it i) does not account for the inherent biases present in the antigen processing and presentation machinery used to generate natural pMHC ligands in cells; ii) interrogates TCR in the context of supraphysiological abundance of pMHC ligand; iii) is indirect and may not be able to identify cross-reactivities which do not have significant sequence similarity to the index peptide or represent distinct epitope clusters^31^; and iv) requires a different custom set of peptide libraries to be generated for the study of each new index peptide. In contrast, we present Tope-seq as a complementary approach for functional cross-reactivity assessment that preserves natural expression, processing, and presentation of library peptides. Genetically encoded cell libraries containing a defined panel of potential epitopes, such as all possible self-antigens, also provide an advantage as they can be directly searched without the need to infer cross-reactivity based on results from degenerate peptide. Additionally, the minigene library system used in conjunction with Tope-seq assay readouts can be flexibly re-used in studies of different TCR, HLA, or target peptide contexts without re-synthesis.

To validate the Tope-seq approach to the assessment of comprehensive self-antigen libraries, it was essential in this study, as in any functional live cell assay, to consider non-specific background from endogenous peptides in the target cell chassis. We found that epitopes of the test TCR derived from naturally expressed host cell proteins were indeed a confounding factor in analyzing minigene library-derived epitopes and we deployed solutions such as CRISPR/Cas9 removal of known expressed epitopes and adaptation of alternative cell chassis lines to address this issue. Though successful in the present study, these solutions are not universal since endogenous background epitopes may not be known or may exist in an indispensable gene. We, therefore, also sought to overcome endogenous peptide background by performing iterative biopanning through minigene libraries, progressively enriching hits on each round. We did this by directly re-cloning and re-expressing primary DNA from captured minigenes for second-round Tope-seq analysis, which was an effective approach for increasing enrichment in known epitopes even in target cell chassis where endogenous epitopes were present.

We also noted enrichment of spurious hits in our WPC and WES self-antigen Tope-seq experiments. This finding was mirrored in other studies of self-antigen assessment using similar contemporaneously developed GZMB based library screening technology where investigators contended with it by performing as many as eight replicate experiments to sufficiently reduce background enough to characterize wild-type, non-affinity-enhanced TCR^32^. Here, we aimed to improve on this by implementing an improved bioinformatic workflow for calling hits from relatively few replicate experiments. We also found that the biopanning approaches we describe above present a feasible technique for further improving signal-to-noise ratios in large library screening campaigns.

The selection of method for minigene library production is another important consideration in the development of Tope-seq screening campaigns. The flexible and modular nature of the Tope-seq system enables a variety of DNA sources to be accessed as a source of minigene library. In our work, we describe two alternative pathways for producing panels of peptide-coding sequences to be interrogated by TCRs-of-interest. In one iteration, we used a defined, computationally-designed, array-synthesized version of a self-antigen library while in the other, genomic DNA from primary samples was sheared and exome-captured. We envision that a variety of minigene library types could be developed and deployed in Tope-seq screening campaigns depending on specific experimental considerations. In Table 3, we contemplate the advantages and disadvantages across the spectrum of minigene library formats.

**Table 3.** Multiple routes are available for constructing genetically encoded minigene libraries to screen by Tope-seq. Two major methods for deriving minigene libraries for Tope-seq analysis were described in this work: synthesized DNA from reference proteome and exome-captured genomic DNA fragments. However, applying the same principles of library construction, screening, and data analysis demonstrated here, other library formats could be contemplated, including cDNA fragment libraries, whole-genome fragment libraries, or synthesized DNA from degenerate, randomized/semi-randomized sequence design. Each of these formats carry specific advantages and disadvantages with respect to the search space afforded, potential gaps or missing sequences inherent, sources of possible background, and issues with DNA sequence fidelity. As a result, we propose a continuum of use cases in which different library construction methods may be optimal for distinct settings in which investigators would seek to test TCR clones or populations against large sets of possible epitopes.

|  | Synthesized DNA from reference proteome | Exome-captured genomic DNA fragments | cDNA fragments | Whole-genome DNA fragments | Randomly generated oligos |
| --- | --- | --- | --- | --- | --- |
| **Complexity** | Low/Medium (costly to scale-up) | Medium | High | Very high | Extremely high (very difficult to comprehensively search) |
| **Gaps** | Polymorphism/strain variation.<br>Non-coding regions. | Polymorphism/strain variation.<br>Exon-exon junctions.<br>Non-coding regions. | Polymorphism/strain variation.<br>Rare transcripts.<br>Tissue-specific transcripts. | Polymorphism/strain variation.<br>Exon-exon junctions. | Theoretically minimal. |
| **Background** | Artifacts from synthesis errors. | Flanking intronic sequence.<br>Out-of-frame peptides. | 5'-/3'-UTRs.<br>Out-of-frame peptides.<br>Highly abundant transcripts. | Non-coding regions.<br>Out-of-frame peptides. | Theoretically maximal. |
| **Fidelity** | Limited by DNA synthesis coupling efficiency (~1 error per $3 \times 10^3$ bp). | High, only limited by PCR error rate. | Limited by reverse transcriptase error rate (~1 error per $2 \times 10^3$ bp). | High, only limited by PCR error rate. | Very high, only limited by biases in coupling efficiency |
| **Advantages** | Expression frame can be strictly controlled.<br>No primary tissue required. | Cost-efficient<br>No up-front sequencing data required. | Cost-efficient<br>No up-front sequencing data required.<br>More coverage than exome-only library. | Cost-efficient<br>No up-front sequencing data required.<br>More coverage than transcriptome-only library. | Cost-efficient (pooled synthesis)<br>No up-front sequencing data required.<br>No primary tissue required.<br>Most unbiased format. |
| **Use cases** | Neo-antigen panels.<br>Microbial ORF libraries.<br>Fine epitope mapping of individual proteins.<br>Self-antigen libraries. | Unbiased primary tumor isolate libraries.<br>Minor histocompatibility antigen discovery<br>Cryptic epitopes<sup>33</sup> | Unbiased primary tumor isolate libraries.<br>Minor histocompatibility antigen discovery<br>Cryptic epitopes<sup>33</sup> | Unbiased primary viral/microbial isolate libraries.<br>Cryptic epitopes<sup>33</sup> | Training data for *in silico* models. |

In conclusion, we have described the latest advances in our Tope-seq methodology, including the development of comprehensive synthesized and captured DNA-coded self-antigen minigene libraries, sAPC chassis cell line engineering, and techniques for iterative functional minigene re-screening. We show that this technology has significant potential utility in assessing functional off-target and dangerous cross-reactivity against healthy tissues in therapeutic TCR lead candidates. We expect the application of function-based high-throughput screening to early discovery in therapeutic TCRs will enable effective de-risking, triage, and re-engineering of lead candidate molecules prior to resource intensive pre-clinical analysis.

## Methods

### Plasmid DNA propagation and isolation

NEB Stable Competent *E. coli* (NEB) was used for the propagation of DNA unless otherwise specified. Bacterial transformations were performed according to manufacturer protocol. *E. coli* was grown in LB broth at 30°C, shaking at 250 rpm. For solid medium, LB broth was supplemented with Bacto agar (1.5% [w/v]; Difco). Media were further supplemented with 100 μg/mL carbenicillin or 50 μg/mL kanamycin as appropriate. Plasmid DNA was isolated using Invitrogen PureLink HiPure Plasmid Filter Maxiprep Kit. All synthesized DNA was produced by Integrated DNA Technologies unless otherwise specified. All cloning enzymes were sourced from New England Biolabs unless otherwise specified. Sanger sequencing verification was performed at all intermediate steps by Genewiz.

### DNA fragment library preparation

WPC design was performed using a custom Python pipeline to produce sets of overlapping non-redundant 60 amino acid fragments (with a minimum 30 amino acid overlap) generated from individual protein sequences from UniProt proteome accession number UP000005640. Extracted amino acid sequences were back-translated using randomized codon usage and designed with Illumina adapter sequences flanking proteome-coding minigenes to facilitate cloning and downstream sequencing. We chose to use non-native codons for producing minigene sequences to mitigate concatemerization of overlapping single-stranded DNA in downstream PCR-based library cloning in plasmid vector. The designed set of DNA sequences (216bp x 535,815 sequences) was synthesized on array-based platforms by Twist Biosciences and delivered as a pool of single-stranded DNA. Library was rendered double stranded and amplified by PCR with Phusion polymerase (1 ng, 30 cycles) prior to being used as input for library cloning procedures. For WES library construction, 1 µg of human genomic DNA (Promega cat. #G3041, lot #0000387521) was shipped to Genewiz for Illumina whole exome-sequencing library preparation only. Prepared exome library was shipped back from Genewiz and PCR-tailed (15 ng, 25 cycles) to amplify library fragments and prepare for cloning into lentiviral transfer plasmid. For WES second-round libraries, 100 ng of genomic DNA sampled from Shifted gate cells captured in first round experiments was subject to PCR amplification and re-cloning. The PCR primers used for all library constructions described here were composed of universal Illumina adapter binding regions tailed with I-SceI/PI-SceI endonuclease sites (F: 5’- TAGGCAAATCTAGGGATAACAGGGTAATTACGACGCTCTTCCGATCT-3’and R: 5’- ATCGATACATCCATTTCATTACCTCTTTCTCCGCACCCGACATAGATCGTGTGCTCTTCCGATCT-3’) and were HPLC- and PAGE-purified (IDT).

### Construction of HLA plasmids

Lentiviral transfer plasmids were derived from pCCL-c-MNDU3-X backbone (Addgene #81071). To produce an acceptor cassette for HLA allele sequences, a custom synthesized cassette (configured as HLA receiver stuffer-P2A-ECFP-GZMB cleavage substrate-EYFP) was inserted by endonuclease cloning between the BamHI/XhoI restriction sites in the original plasmid (BamHI site ablated during this step) to yield the pHLAI-FRET backbone. Coding sequence for the HLA-A*01:01 allele-of-interest was custom synthesized and cloned into designated HLAI positions via NheI/MluI restriction cloning (restriction sites introduced in acceptor plasmid construction).

### Construction of minigene plasmids

Lentiviral transfer plasmids were derived from pCCL-c-MNDU3-X backbone (Addgene #81071). To produce an acceptor cassette for individual minigenes or minigene libraries, a custom synthesized cassette (configured as Minigene receiver stuffer-IRES-mStrawberry) was inserted by endonuclease cloning between the BamHI/XhoI restriction sites in the original plasmid (BamHI site ablated during this step) to yield the pMGI backbone. Synthesized minigenes and captured DNA fragment minigenes were PCR amplified with primers annealing to proximal adapter sequence flanking minigene fragments and tailed with I-SceI/PI-SceI meganuclease recognition sites. Individual minigenes were cloned into pMGI by standard I-SceI/PI-SceI restriction cloning. Minigene libraries were cloned into pMGI using a specialized multistep digestion/ligation procedure to increase library cloning efficiency and fidelity. Briefly, this procedure consists of i) PI-SceI/PmeI-linearized pMGI ligated under high DNA concentration T4 ligation conditions with PI-SceI-treated library fragments, ii) secondary digestion of linear pMGI/library intermediate with I-SceI, and iii) second low-concentration ligation of I-SceI treated linear pMGI-library plasmid to re-circularize final library vector. Prepared pMGI-library circular plasmids were bacterially expanded by transformation of MegaX DH10B T1^R^ *E. coli* (ThermoFisher) by electroporation (Bio-rad GenePulser II) according to manufacturer protocol and 18-hour, 37°C outgrowth on 576 cm^2^ solid agar surface prior to colony scraping and plasmid isolation. Further detail on high-throughput cloning protocol can be found in Appendix A of ref^34^.

### Construction of TCR plasmids

Lentiviral transfer plasmid was derived from the pCCL-c-MNDU3-X backbone (Addgene #81071). To produce an acceptor cassette for TCR, a custom synthesized cassette (configured as a TCR receiver stuffer-P2A-mStrawberry) was inserted by endonuclease cloning between the BamHI/XhoI restriction sites in the original plasmid (BamHI site ablated during this step) to yield the pTCR backbone. EB81.103 and a3a TCR cassettes were synthesized as TCRɑ-T2A-TCRβ fragments and were cloned into pTCR via BamHI/EcoRI restriction cloning (restriction sites introduced in acceptor plasmid).

### Mammalian cell culture

All cell cultures were maintained in RPMI-1640 supplemented with 2 mM GlutaMAX, 1 mM sodium pyruvate, 50 μM β-mercaptoethanol, 10 mM HEPES, 100 U/mL penicillin, 100 U/mL streptomycin, 1X MycoZap™ Prophylactic (Lonza) and 10% heat-inactivated fetal bovine serum. Culture media and supplements were all sourced from Gibco unless otherwise indicated. K562 and HEK-293T were sourced from ATCC; 721.221 cells were a kind gift from the Judy Lieberman lab (Boston Children’s Hospital). All cell cultures were maintained at 37°C and 5% CO2 atmosphere. K562 based cell lines were started at 2x10^5^ cells/mL and subcultured every 4 days by diluting cells 1/10. HEK-293T were passaged every 4 days by trypsinizing, washing 1X with PBS, and re-seeding cells at 4x10^6^ cells/flask in T-75 format. 721.221 cell lines were started at 1x10^5^ cells/mL and subcultured every 3 days by diluting cells 1/10. Cell counts were performed using an EVE automated cell counter. All cell lines were certified to be free of mycoplasma contamination using the Venor GeM Mycoplasma Detection Kit (Sigma) prior to experimental data collection. All mammalian cell cultures were incubated at 37°C and 5% CO2.

### Virus production

Lentiviral vectors were produced by combining 180 µg of each transfer plasmid with 162 µg of pCMV-ΔR8.91 and 18 µg of pCMV-VSV-G plasmids. These DNA mixes were incubated with 18 mL OptiMEM and 1 mL of TransIT-LT1 reagent (Mirus) for 30 minutes at room temperature. To 12 x T-75 culture flasks containing between 16 – 24 million HEK-293T cells, old media was removed and replaced with 10 mL fresh, pre-warmed media and 1.5 mL of transfection mix per plate. Media was again replaced 18 hours post-transfection with 10 mL of pre-warmed fresh media. Viral supernatant was then collected at 48 hours post transfection (replacing with 10 mL pre-warmed fresh media) and 72 hours post-transfection. To concentrate virus, pooled supernatants were ultracentrifuged (100,000 RCF, 90 minutes, 4°C). Viral pellets were resuspended at 4°C overnight in 1 mL OptiMEM. Titers of viruses were determined by testing 2, 4, 8, 16, 32 or 64 μL of 10X diluted virus on 1x10^5^ K562 cell/well in a final volume of 500 μL of media in 24-well format. Transduction efficiency was determined by measuring the % of fluorescent cells detected in flow cytometry. Values were used to determine K562-infecting units (KIU) per µL of undiluted virus concentrate.

### TCR-T cell production

Peripheral blood leukocytes used as a source of primary T cells were obtained from Stemcell Technologies. We report, in line with BRISQ Tier 1 recommendations that human biospecimens used in this study were leukapheresis products isolated from live normal human donors. Volunteer donors were screened for complete blood count, serum proteins, hematocrit, and confirmed negative for HIV-1, HIV-2, hepatitis B and C prior to procedure. Blood was drawn directly from arm vein into leukapheresis machine, and the isolated leukocyte fraction was collected directly into sample bag containing acid citrate dextrose solution A anticoagulant and stored/shipped at 0-4°C. On receipt, leukopaks were processed using a Ficoll-Hypaque density gradient centrifugation and ACK buffer red-blood cell lysis procedure, both according to manufacturer provided protocols.

The resulting isolated PBMCs were aliquoted into individual vials of 1x10^7^ cells and cryopreserved in liquid nitrogen vapor phase in heat-inactivated fetal bovine serum + 10% DMSO. Prior to each Tope-seq experiment, ten vials of PBMC (∼10^8^ cells total) were thawed and FACS-sorted to isolate CD8^+^ CD4^-^ CD56^-^ cells (using BV785-conjugated anti-CD8 antibody, clone SK1 (Biolegend); PerCP-Cy5.5-conjugated anti-CD4 antibody, clone RPA-T4 (Biolegend); and FITC-conjugated anti-CD56 antibody, clone HCD56 (Stemcell)) and rested for 48 hours in complete RPMI + 300 IU/mL human IL-2 (Stemcell) at a cell density of 10x10^6^ cells/mL. Simultaneous T-cell transduction and activation was performed by counting CD8^+^ T cells after the rest period, combining cells with 30 KIU/live cell of TCR-encoding lentiviral vector, and transferring the cell/lentivirus mixture to wells of a non-TC treated, flat-bottom 96-well plate pre-coated with LEAF-purified anti-CD3 antibody, clone OKT3, and anti-CD28 antibody, clone 28.2 (eBioscience). Cells were stimulated for 20-24 hours on coated wells at 37°C and 5% CO2 atmosphere. After the transduction/stimulation period, cells were removed from coated wells, diluted to 1x10^3^ cells/mL in complete RPMI + 300 IU/mL hIL-2, and plated in U-bottom 96-well plates. Media was 50% changed every 4 days and used in co-culture experiments 11-14 days after stimulation.

### Creation of HLA-FRET sAPC lines

K562 cell lines were transduced with viral vector produced using pHLAI-A0101-P2A-FRET plasmid at an MOI of 1 KIU/cell. 721.221 cell lines were transduced with viral vector produced using pHLAI-A0101-P2A-FRET plasmid at an MOI of 3 KIU/cell in serum-free OptiMEM for 24 hours prior to returning cells to complete media. Transductions were performed by placing 1x10^5^ cells/well in a final volume of 500 μL of media in a single well of 24-well format for 48 hours before resuming normal culture technique to expand. Purified HLA-expressing K562 and 721.221 (KFRET.A0101 and 721FRET.A0101, respectively) were isolated by FACS to recover strong FRET expressers and HLA surface expression (based on staining with APC-conjugated anti-human MHC, clone G46-2.6, BD Bioscience).

### Creation of single-minigene/minigene library-expressing sAPC lines

Minigene-expressing target cells were produced by transducing pre-purified KFRET.A0101 or 721FRET.A0101 with viral vector produced using pMinigene-IRES-mStrawberry format constructs. For single-minigene controls (i.e. MAGEA3 and TTN minigene-expressing lines), KFRET.A0101 were transduced at an MOI of 0.3 KIU/cell and 721FRET.A0101 was transduced at an MOI of 1 KIU/cell under the same scale and conditions as HLA-FRET cell line creations. For WPC minigene libraries in KFRET.A0101, cells were transduced with library vector at MOI = 0.2 over 3.5 x 10^8^ starting cells (in G-Rex format) for low-MOI approach or MOI = 7 over 1.0 x 10^7^ starting cells for high-MOI approaches. For 721FRET.A0101 library transduction, 8 x 10^6^ cells were infected with 20 KIU/cell in serum-free OptiMEM (to prevent cluster formation during viral infection). After 24 hours, transduced 721FRET.A0101 were centrifuged and re-seeded in complete medium for continued culture. For 1^st^ round WES minigene libraries in KFRET.A0101, cells were transduced with library vector at MOI = 3 over 8 x 10^6^ starting cells and, in 2^nd^ round panning libraries, at MOI = 0.7 over 1.2 x 10^7^ starting cells. For all minigene-expressing cell lines, purity sorting was conducted by selecting RFP^+^ cells. Minigene library-expressing cell lines were purity-sorted 5-7 days after transduction (yielding a minimum of 1.5 x 10^7^ founder clones for WPC format; 2.5 x 10^7^ for WES/panning format) and cryopreserved 5-7 days after purity sorting to minimize library drift (stored at 2.5x10^7^ cells/vial to mitigate bottlenecking).

### Co-culture setup for Tope-seq experiments

For large-scale library screening Tope-seq experiments, TCR-T cells were always freshly prepared starting 11-14 days prior from primary human PBMC, while library-expressing sAPC were always thawed 5 days prior to assay setup using an inoculum of >2.5x10^7^ cells in a 300 mL media volume. Replicate experiments were always performed on different days, using TCR-T cells produced from different donor sources, and library target cell lines prepared from separate transductions. To prepare experimental co-cultures, target and effector cells were adjusted to a density of 2 x 10^6^ cells/mL and 3 x 10^6^ cells/mL, respectively in fresh, pre-warmed media (TCR-T were adjusted to a higher density to account for an average 65% TCR transduction rate). For KFRET based library cell lines, 50 mL of each were combined and gently swirled to mix. Cell mixture was distributed uniformly to all wells of 10 x 96-well U-bottom plates and incubated for 12 hours. For library screens involving 721FRET based targets, where very short (<1hr) co-culture periods were needed, plates were prepared in staggered batches: target and effector cells were each adjusted to a density of 2 x 10^6^ cells/mL or 3 x 10^6^ cells/mL, respectively, in fresh, pre-warmed media and 15 mL of each were combined in a 50 mL centrifuge tube. Cell mixture was gently inverted to mix and distributed uniformly to all wells of 3 x 96-well U-bottom plates. During sorting of first batch, the next batch was assembled, incubated, and sample prepped. Once complete, new batch was immediately loaded for sorting. For each replicate, three staggered batches were prepared and run. In all screening experiments, a plate of individual control co-cultures was assembled, consisting of TCR-T cells against single-minigene and no-minigene controls. This was done by adjusting each target and effector population to a density of 2 x 10^6^ cells/mL in fresh, pre-warmed media, combining 100 μL of each test pair to run, and plating in 2 wells of a 96-well U-bottom plate per control condition to be included. For each control condition, a pair of wells was left unmixed and combined immediately before flow cytometry to serve as a T0 loading control. On conclusion of co-culture period, cells were stained on-plate with 0.5% BV785-conjugated anti-CD8 antibody, clone SK1 (Biolegend) and 0.1% Fixable Viability Dye eFluor 780 (ThermoFisher) for 15 minutes at 4°C before transferring co-culture wells to tubes, washing cells 1X in 10 volumes cold PBS, and resuspending in cold PBS + 2% FBS. Prepared co-culture samples were kept on ice and taken immediately for flow cytometry/FACS.

### Flow cytometry/FACS

All FACS was performed BD FACSAria Fusion running BD FACSDiva v9.0 software. Gating was performed by monitoring FVD eFluor780 channel (ex. 640, em. 780/60), RFP channel (ex. 561, em 610/20 + 600LP), YFP channel (ex. 488, em. 530/30 + 505LP), PerCP-Cy5.5 channel (ex. 488/em. 695/40 + 655LP), BV785 channel (ex. 405, em. 780/60 + 750LP), FRET channel (ex. 405, em. 525/50 + 505LP), and/or CFP channel (ex. 405, em. 450/50) according to experiment design. Cytometric analyses were performed on BD LSRFortessa and quantitated using BD FlowJo v10.

### Minigene sequencing

Cells sorted from matched FRET-shifted and unshifted gates in each replicate Tope-seq screening experiment were immediately pelleted and lysed using DNAzol reagent. Ethanol precipitation, washing, and re-solubilization was performed according to manufacturer’s protocol. To prepare samples for sequencing, two rounds of PCR were conducted. In the first, 50% of all recovered gDNA per sample was input as template across multiple parallel reactions (capping the amount of template to be added per 100 µL reaction at the lower of 10 µL or 250 ng). First round PCR was run for 25 cycles with Phusion polymerase (NEB) and Illumina adapter primers, F: 5’- ACGACGCTCTTCCGATCT-3’ and R: 5’-CGTGTGCTCTTCCGATCT-3’. For second round PCR, 100 ng of column-cleaned first round product was used as template and amplified for 5 cycles using primers from NEBNext Multiplex Oligo Set, uniquely barcoding each gate sample of each replicate experiment. Prepared libraries were subject to PE150 sequencing on the Illumina HiSeq 4000 platform at off-site service provider, Genewiz.

### Data analysis

For WPC library-based screening, matched Shifted and Unshifted gate minigene sequencing reads were trimmed with Cutadapt v1.18 (default settings), paired-end merged with FLASh v1.2.11 (--min-overlap 20 -- max-overlap 150 --max-mismatch-density 0.2 --allow-outies), quality filtered with FASTX-toolkit (keeping reads with 90% of bases > Q20), and error corrected/mapped to synthesized minigenes using Starcode v1.1 (-d 4 in spheres mode for collapsing errors; -d 3 in spheres mode for aligning to canonical minigene designs). Motif-building analyses were conducted using NetMHC4.0 (default parameters) and GibbsCluster2.0 (lambda 0, sigma 0, trash cluster threshold 10). For WES libraries, Illumina adapters were removed using Trim Galore v.0.6.6 in paired-end mode. A quality score of 20 was used for trimming low quality bases from the read ends. Reads were aligned to the hg19 human reference genome using BWA-MEM2 v.2.2.1. Reads were then processed using samtools v.1.9 by name ordering and collating, fixing mate information, position ordering, marking duplicates and indexing. Processed reads were counted by using a sliding reference window to count all in-frame minigene fragments traversing each peptide-coding window of defined size in the entire coding genome. Data handling and visualization for all sequencing experiments was performed with R v4.4.1, seqinr v4.2-36, and ggplot2 v3.5.1.

## Declarations

### Ethics approval

The study protocol was approved by the BC Cancer Research Ethics Board (REB) in accordance with the guidelines set out in the Declaration of Helsinki. All human samples used in this study came from commercial vendors; no donor material was isolated at the BC Cancer Research Institute. Use of commercially obtained human materials at the BC Cancer Research Institute was done under the oversight of the BC Cancer REB. Human primary T cells used in this study were obtained from Stemcell Technologies and were derived from leukopaks collected in the U.S. from donors providing written informed consent and under the approval of Hummingbird IRB. Human genomic DNA used in this study was obtained from Promega and derived from whole blood collected in the U.S. from donors providing written informed consent and under the approval of Promega’s Human Subjects Review Board.

### Availability of data

Raw flow cytometry data generated in this study are available from the lead or corresponding author on reasonable request. Raw fastq files and processed minigene counts from library QC and Tope-seq experiments are available for download from the NCBI Sequence Read Archive under BioProject accession number PRJNA1258577.

### Code availability

Custom code used in this study may be accessed from https://github.com/DrGovindaSharma.

### Author contributions

GS developed the experimental strategy with feedback from FT, JR, RAH. *In silico* design of hWPC minigene library was conducted by SS and SDB with guidance from GS. Molecular cloning of TCR, HLA, and minigene library plasmids was conducted by GS and JR. Viral vectors and transduced cell lines were produced by GS and FT. Cell screening for all experiments was conducted by GS and FT. Minigene sequencing library preparation for Tope-seq experiments were performed by GS and JR. Data analysis was performed by GS and SAS. Manuscript preparation was done by GS with editing and feedback from SDB, SAS, FT, and RAH.

## Acknowledgements

The authors would like to acknowledge Dr. Vikram Juneja and Dr. Kendra Foley for their valuable input and ideas.

