## Supplementary Tables and Figures for "Comprehensive Functional Self-Antigen Screening to Assess Cross-Reactivity in a Promiscuous Engineered T-cell Receptor"

| **Supplementary Table 1. Summary of WPC plasmid library quality assessment by NGS.** A sample of WPC minigene library plasmid was input for Illumina library preparation and sequencing to quantify non-mappable reads and design coverage. | |
|---|---|
| | **Count** |
| Quality filtered reads | 169,864,738 |
| Mapped to reference | 165,969,580 (97.7% of total quality filtered reads) |
| Reference coverage | 522,045 (97.4% of designed reference library) |

**Supplementary Table 2. Summary of low-MOI KFRET.A0101.WPC quality assessment by NGS.** A sample of WPC library-expressing target cells generated by initial low MOI strategy were taken from each of four different replicate transductions. For each sample, 2-3 x $10^7$ cells were lysed and subject to minigene recovery by PCR from isolated gDNA prior to Illumina sequencing. For each individual replicate, coverage of designed library and average read depth for each minigene was quantitated. Reads across all four replicates were also merged and used to quantify the design library coverage as a union dataset.

| | **Replicate 1** | **Replicate 2** | **Replicate 3** | **Replicate 4** | **Union** |
|---|---|---|---|---|---|
| Quality filtered, mapped reads | 247,448,584 | 193,723,485 | 126,096,987 | 125,836,195 | |
| Reference coverage (count) | 416,474 | 473,256 | 486,634 | 487,257 | 518,993 |
| Reference coverage (%) | 77.7 | 88.3 | 90.8 | 90.9 | 96.9 |
| Average read depth per minigene | 594.1 | 409.3 | 259.1 | 258.3 | |

**Supplementary Table 3. Summary of high-MOI KFRET.A0101.WPC quality assessment by NGS.** A sample of WPC library-expressing target cells generated by high MOI strategy were taken from each of two different replicate transductions. For each sample, 2-3 x $10^7$ cells were lysed and subject to minigene recovery by PCR from isolated gDNA. For each individual replicate, coverage of designed library and average read depth for each minigene was quantitated.

| | **Replicate 1** | **Replicate 2** |
|---|---|---|
| Quality filtered, mapped reads | 176,219,645 | 178,408,919 |
| Reference coverage (count) | 524,078 | 522,862 |
| Reference coverage (%) | 97.8 | 97.6 |
| Average read depth per minigene | 336.2 | 341.2 |

**Supplementary Table 4. Summary of KFRET.A0101.WES quality assessment by NGS.** A sample of 2 x $10^7$ cells were lysed and subject to minigene recovery by PCR from isolated gDNA. Raw reads were quality filtered and mapped to hg19 human reference genome. The WES library was estimated to encode 100X coverage of the human exome (based on number of unique mapped reads detected, average fragment length, and an estimated exome size of 50 Mbp). The average read depth of individual minigenes was also quantified and found to be much lower than that typically observed in WPC libraries, indicating that epitope redundancy within libraries of WES format is derived primarily from high unique minigene fragment count.

| | |
|---|---|
| Raw reads | 114,717,984 |
| Quality filtered, mapped reads | 108,698,439 (94.8% of total reads) |
| Average fragment size (bp) | 212 |
| Num. unique fragments | 25,205,103 |
| Estimated exome coverage | >100X |
| Average read depth per minigene | 4.3 |

**Supplementary Table 5. Quantification of WES library bottlenecking in 2$^{nd}$ round libraries.** The number of unique minigenes detected in 2$^{nd}$ round screening of a3a and EB81.103 TCR-T cells after re-cloning Shifted gate minigenes from initial 1$^{st}$ round screens. In both cases, a substantial fraction of minigenes from the original source WES minigene library are not re-detected in 2$^{nd}$ round experiments.

| | **a3a-screened 2$^{nd}$ round library** | **EB81.103-screened 2$^{nd}$ round library** |
|---|---|---|
| Quality filtered, mapped reads | 63,386,248 | 185,812,985 |
| Num. unique fragments | 1,043,412 | 1,935,310 |
| Percentage of original library | 4.1% | 7.7% |

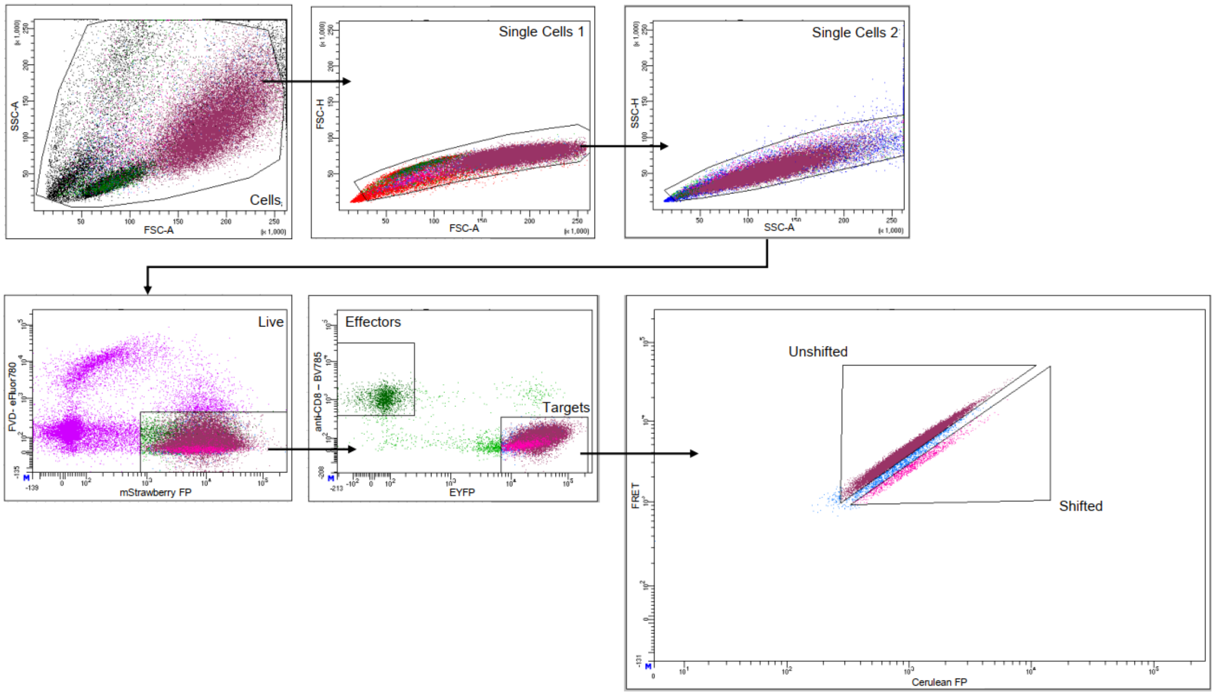


**Supplementary Figure 1. Representative gating strategy for FRET-shift flow cytometry in Tope-seq experiments.** A common sorting template was used to perform two-way FACS for capturing matched Shifted and Unshifted gate cells in all library screens used to generate Tope-seq data. Intact, single cells are first selected using sequential forward- and side-scatter gates. Viable cells are then selected by gating on cells negative for FVD780 vital staining dye and positive for mStrawberry RFP living color. Effector and target cells are separated by CD8-BV785 staining and EYFP expression. Unshifted and Shifted gates are drawn by adjusting gate boundaries to keep background FRET-shift in $T_0$ or targets-only control samples to <1%.

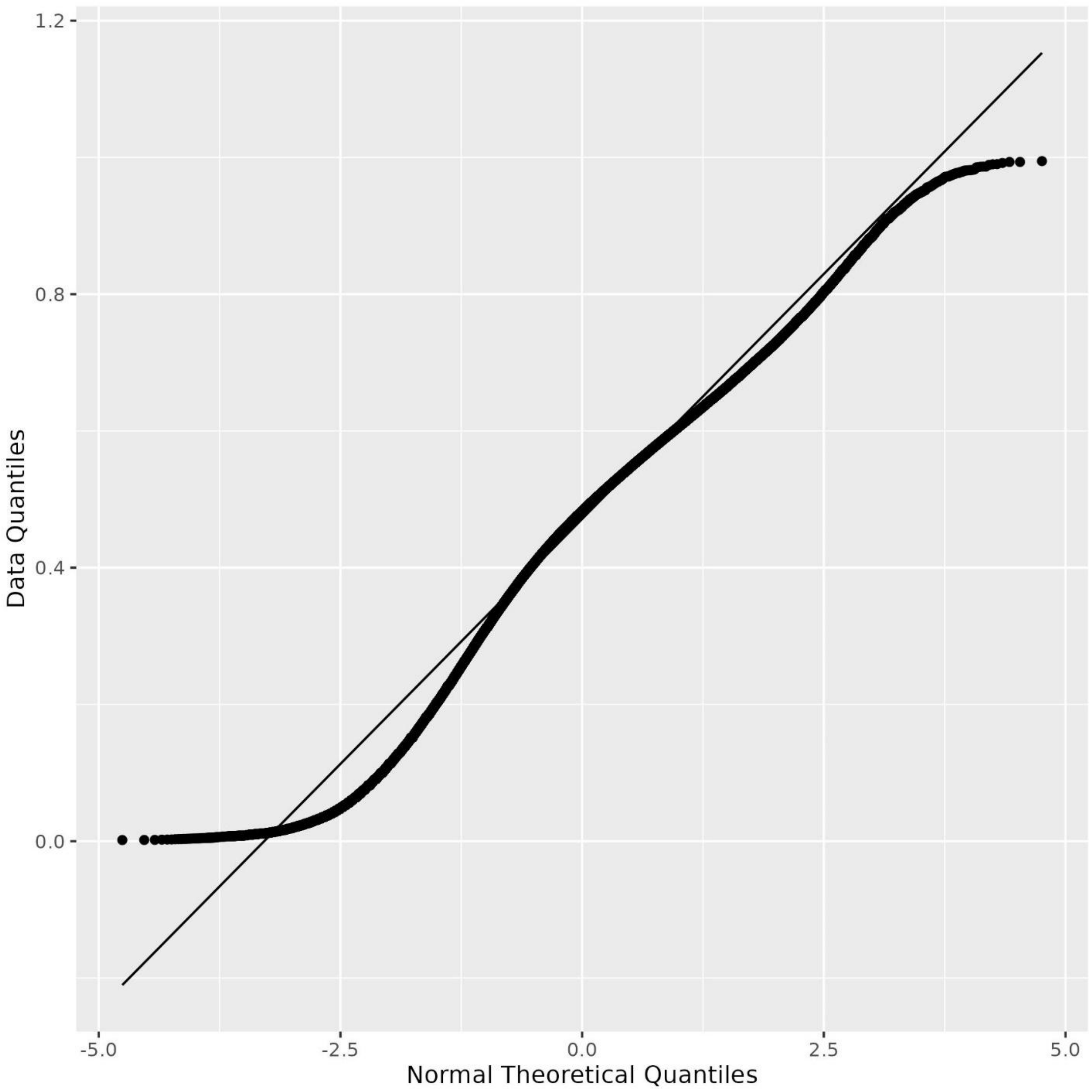


**Supplementary Figure 2. Quantile-quantile plot to estimate normality of Tope-seq scores.** To confirm if Z-score calculation was suitable for significance, normality testing was done on enrichment scores calculated from relative frequency of normalized read count for each minigene/peptide sequence in Shifted gate as a proportion of its normalized read counts in Shifted + Unshifted. Strong linear correlation in a representative example, shown here, between theoretical quantiles and experimental quantiles empirically confirmed that the relative frequency scoring statistic used in this study are approximately normally distributed.

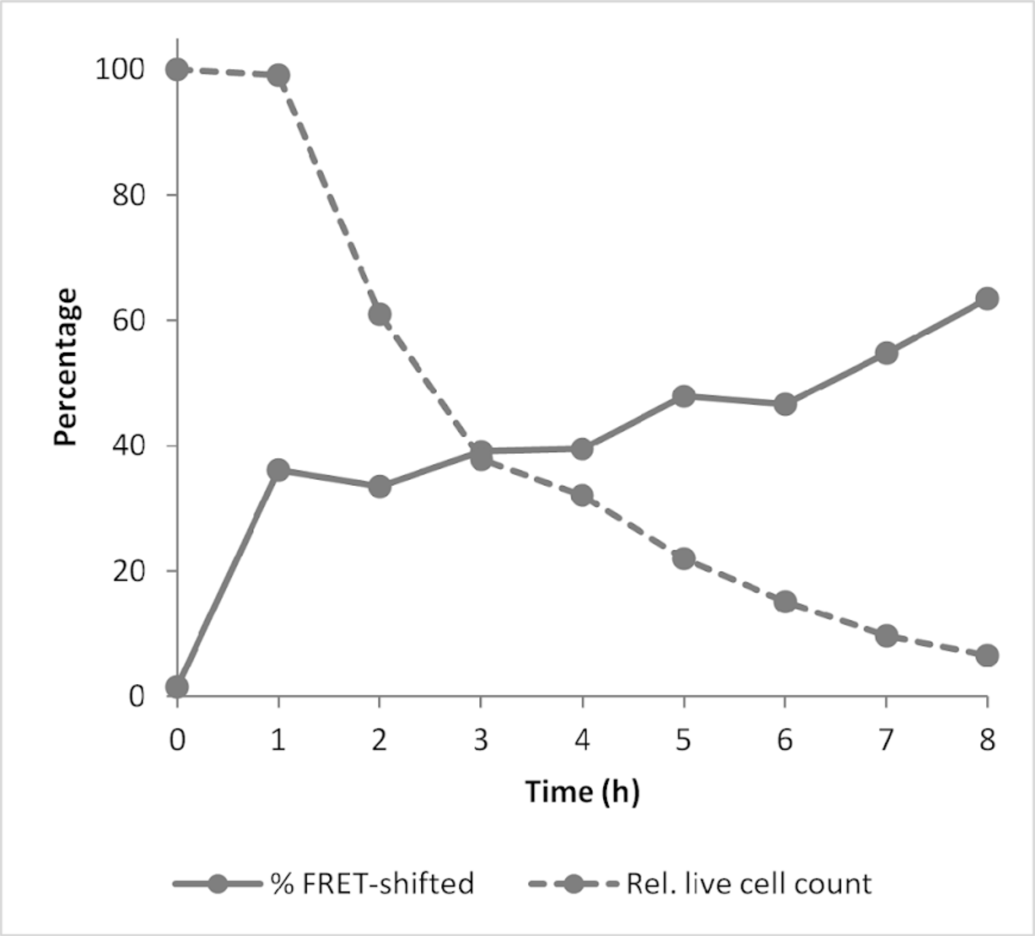


**Supplementary Figure 3. Time course assessment of a3a TCR-T + 721FRET.A0101.MAGEA3$^{143\text{-}202}$ co-cultures.** To validate 721.221 cell line for use as a target sAPC to use in Tope-seq experiments, a time course experiment was performed to determine the safe-sorting window within which GZMB-loaded target cells should be collected before being lost to apoptosis cell death and degradation. Engineered 721.221 cells expressing HLA-A*01:01/GZMB FRET reporter transgenes and a positive control MAGEA3$^{143\text{-}202}$ minigene cassette were prepared by lentiviral transduction and used for time-course testing. Reactions were assembled by plating $2 \times 10^5$ of each cell line into 2 separate wells of a U-bottom 96-well plate. An effector/target well pair was prepared for each of the 8 time points investigated and left unmixed until co-culture periods were initiated at the appropriate time prior to flow cytometry by combining effector and target wells and re-distributing co-cultures to both wells. Flow cytometry was conducted on all samples in a single batch at experiment conclusion. Percent FRET-shift signal and percent survival (live target cell counts normalized to matched $T_0$ control) are displayed.
